# Thyroid hormone regulates abrupt skin morphogenesis during zebrafish postembryonic development

**DOI:** 10.1101/2021.04.09.439217

**Authors:** Andrew J. Aman, Margaret Kim, Lauren M. Saunders, David M. Parichy

## Abstract

Thyroid hormone is a key regulator of post-embryonic vertebrate development. Skin is a biomedically important thyroid hormone target organ, but the cellular and molecular mechanisms underlying skin pathologies associated with thyroid dysfunction remain obscure. The transparent skin of zebrafish is an accessible model system for studying vertebrate skin development. During post-embryonic development of the zebrafish, scales emerge in the skin from a hexagonally patterned array of dermal papillae, like other vertebrate skin appendages such as feathers and hair follicles. We show here that thyroid hormone regulates the rate of post-embryonic dermal development through interaction with nuclear hormone receptors. This couples skin development with body growth to generate a well ordered array of correctly proportioned scales. This work extends our knowledge of thyroid hormone actions on skin by providing in-vivo evidence that thyroid hormone regulates multiple aspects of dermal development.

**Highlights:**

- Thyroid hormone (TH) is necessary for normal squamation patterning in zebrafish.
- Stratified dermis develops by migration of primary hypodermal cells.
- Dermis stratifies in an invariant wave.
- TH regulates the rates of multiple aspects of dermis development.
- Scale size and density are sensitive to skin size at onset of squamation.

## 1. Introduction

From the spectacular metamorphosis of jellyfish and frogs to the explosive growth of long bones in adolescent humans, the adult form of many animals is shaped by extensive postembryonic morphogenesis. In vertebrates, thyroid hormone (TH) is a key regulator of postembryonic development (Laudet, 2011; McMenamin and Parichy, 2013; Tata, 1998). TH is necessary and sufficient for all aspects of amphibian metamorphosis (Tata, 1998) and has been shown to regulate development of the adult pigment pattern, pancreas, skeleton and nervous system in the zebrafish (Hu et al., 2020; Hu et al., 2019; Hur et al., 2017; Matsuda et al., 2017; McMenamin et al., 2014; Saunders et al., 2019; Volkov et al., 2020). One of the many important TH target organs in human is the skin. Defects in thyroid function or peripheral TH signaling are associated with a variety of debilitating skin disorders yet the cellular and molecular pathophysiology of these conditions downstream of TH remains obscure (Mancino et al., 2021).

Skin is a large, distributed organ system that is coupled to, and connected with, the underlying musculoskeletal system. Vertebrate skin is endowed with specialized appendages, commonly referred to as dermal appendages or epidermal appendages, and referred to here simply as “skin appendages.” These include hairs, feathers, eccrine glands, scales and others (Lai and Chuong, 2016). Mechanisms underlying skin appendage patterning and morphogenesis include integrated signaling networks involving Wnt/β-catenin, Fgf, HH, and Eda/Edar/NFk-B signaling pathways, and these are conserved across vertebrates, including zebrafish (Aman et al., 2018; Harris et al., 2008; Iwasaki et al., 2018; Rohner et al., 2009; Sire and Akimenko, 2004).

Zebrafish skin is unusually accessible to experimental manipulation and live imaging and is therefore advantageous for studying organ system development and regeneration (Aman and Parichy, 2020; Cox et al., 2018; De Simone et al., 2021; Rasmussen et al., 2018). The dominant feature of adult zebrafish skin, aside from the magnificent stripes, is an array of ossified scales embedded in the dermis. Scales form relatively late in development and are the penultimate adult trait to form prior to sexual maturity (Parichy et al., 2009; Sire et al., 1997). Here we investigate the role of thyroid hormone (TH) in zebrafish skin development. We show that TH regulates the timing of dermal morphogenesis and we describe essential features of this morphogenesis for the first time. We additionally find that TH couples the precise hexagonal patterning of the scales to the musculoskeletal system by modulating the rate of skin development relative to body growth. Our results provide new insights into TH regulation of postembryonic development and may provide clues to TH-associated pathologies of human skin.

## 2. Materials and Methods

### 2.1. Fish

Fish were maintained in the WT(ABb) background at 28.5°C. Lines used were: *Tg(sp7:EGFP)^b1212^* abbreviated *sp7:EGFP* (DeLaurier et al., 2010);*Tg(7xTCF-Xla.Siam:nlsmCherry)^ia5^* abbreviated *7xTCF:mcherry* (Moro et al., 2012); *Et(krt4:EGFP)^sqet37^* abbreviated *ET37:EGFP* (Parinov et al., 2004); *Tg(ubi:Zebrabow-S^a132^)* abbreviated *ubb:RFP; Tg(tg:nVenus-v2a-nfnB)* abbreviated *tg:Venus-NTR* (McMenamin et al., 2014); *vizinni / gh1^wp22e1^* (vizzini) (McMenamin et al., 2013); *TgBAC(jam3b:jam3b-mCherry)^vp36rTg^* abbreviated jam3b:jam3b-mCherry (Eom et al., 2021); *Tg(6xTETRE-Ocu.Hbb2:EGFP)^bns91Tg^* abbreviated TRE:EGFP (Matsuda et al., 2017); *Tg(ol_osx4.1:nls-EOS)* abbreviated *sp7:nucEOS* (this study). Staging was performed using the Standardized Standard Length system or standard length measurements (Parichy et al., 2009) as estimated in FIJI (Schindelin et al., 2012). Developmental age, in days post fertilization (dpf), is provided as a reference tempo.

### 2.2. In-situ hybridization

All in-situ probe templates were amplified using Primestar-GXL (Takara) from cDNA prepared with SSIII (ThermoFisher) with the following primers: *slc16a10* 5’-tctgtctgatcatcacgctcct, 5’-tctgtctgatcatcacgctcct; *slco1c1* 5’-AACACACTCGTTTTGGACCACA-3’, 5’-GTTGCCTATTTCAAAGCTGCCG-3’; *thraa* 5’-TTTCGCGTTGTTTTGGAAGCAG, 5’-CAGCGGTAATGATAGCCAGTGG-3’; *thrab* 5’-CTCACATGATTGGCTGCTGGAT-3’, 5’-CAGCGGTAATGATAGCCAGTGG-3’; *thrb* 5’-AGCACGAGATATCAGCGAGTCT-3’, 5’-TTGAAGCGACATTCTTGGCACT-3’; rxrab 5’-GAAGATTCTGGAGGCCGAACTG-3’, 5’-CTCGTTCCCTTAATGCCTCCAC-3’; rxrba 5’-AGACCGCAGTGTATCATCAGGA-3’, 5’-TTCCTCACAGTGCGCTTGAAAA-3’; rxrbb 5’-TCGGCGTTTGTTGGAAGATAATCA-3’, 5’-GTCCCCACAAATTGCACACATG-3’ rxrga 5’-CTCTATGCCCACCACTTCCAAC-3’; 5’CCACCGGCATCTCTTCATTGAA-3’; rxrgb 5’-CTCTCAGTTCGCCCTCCATTTC-3’, 5’-TGTGTACGTCTCAGTCTTGGGT-3’. A T7 RNA polymerase binding site with short 5’-tail (aaaaTAATACGACTCACTATAG) was added to reverse primers for transcription using T7 RNA polymerase incorporating DIG labelled ribonucleotides (NEB, Roche). PCR amplified probe templates were confirmed by sanger sequencing.

*In-situ* probes and tissue were prepared as described previously (Quigley et al., 2004). Hybridization and post-hybridization washes using a BioLane HTI 16Vx (Intavis Bioanalytical Instruments) were performed as previously described (Aman et al., 2018).

### 2.3. Transgenic and line production

Generation of *sp7*:nucEOS [Tg(*ol_osx4.1:nls-EOS*)] transgenic line. nucEOS coding sequence was amplified with 5’-aaaaaaGGCGCGCC CACCATGGCTCCAAAGAAGAAG-3’, 5’-aaaaaaGCTGCAGG CTATCGTCTGGCATTGTCAGGC-3’; digested with AscI and SbfI (NEB) and cloned into similarly digested pNVT vector (Renfer et al., 2009) to yield pNVT-nucEOS. Subsequently, a 4.1 kb enhancer of the previously characterized Medaka (Oryzias latipes) sp7(osx) gene (Renn and Winkler, 2009), was amplified using primers 5’-aaaaaaGCGATCGC TGAACATGTCAGTGCCATCAGAATG-3’, 5’-aaaaaaGGCGCGCC CGGGACAGTTTGGAAGAAGTCG-3’, digested with AsiSI and AscI (NEB) and ligated into similarly digested pNVT-nucEOS. The resulting plasmid was injected into zebrafish zygotes together with IsceI meganuclease (NEB) as previously described (Pyati et al., 2005). Injected F0s were raised and and outcrossed to WT(ABb) to establish the line.

*dio2^vp42rc1^* and *tbx6^vp43rc1^* mutant lines were produced by injecting in-vitro transcribed single guide RNAs and recombinant Cas9 protein (PNA Bio) as previously described (Saunders et al., 2019; Shah et al., 2015). The following guide RNA sequences were selected using the CRISPOR webtool (Concordet and Haeussler, 2018): *dio2*, 5’-GCTGTATGACTCGATAGTGC; *tbx6*, 5’-TGAGGCTTTAGCAGTGCAGG-3’. Injected F0s were outcrossed to WT(ABb) and progeny were genotyped by direct Sanger sequencing of PCR products using the following primers: *dio2*, 5’-CAACGGGTAGCCTCTGGTTA-3’; 5’-CGTCTGAGTCACTCCACAGT-3’. *tbx6*, 5’-TTTACTGAAAACCCCATTCAAAAA-3’, 5’-TGATTTATTCACCCTCAAACCATT-3’. All analyses were performed at least three generations following mutagenesis.

### 2.4. Histology

Tissue was fixed in freshly prepared 4% PFA/PBS (EMS) for 1 h at room temperature, equilibrated into 5% gelatin (300-bloom, type-A, Sigma), and sectioned on a Vibratome 1500 (Harvard Apparatus). Sections were stained with CellBright Red (Biotium) diluted 1:500 in PBS for 30 minutes at room temperature, washed 6×10 minutes with PBS and imaged immediately.

### 2.5. Imaging and image processing

Live imaging: larvae were immobilized using 0.2% Tricane Methanesulfonate (Western Chemical) and imaged on an inverted Zeiss AxioObserver microscope equipped with Yokogawa CSU-X1M5000 laser spinning disk or Zeiss LSM880 with Airyscan. Whole fish images were assembled from stitched tiles using Zeiss Zen software. Orthogonal projections were rendered using FIJI (Schindelin et al., 2012). Backgrounds in Movie 1,2; Fig, 1,2,3,4,5,8A, and 9B were masked to remove reflection artifacts. LUT gamma was adjusted in Fig. 8A to highlight migratory dermal cells and Fig. 9B to highlight myosepta. In-situs were imaged on a Zeiss AxioObserver microscope. Brightness and contrast were adjusted using Adobe Photoshop when necessary. Images in Figs. 1A, 2A, 3A, 4A, 5A, 8A, S1A, S10 and in Movie 1 were size normalized for display based on the posterior opercle, pelvic fin base and caudal fin base. Scale bars reflect magnitude of image size scaling.

**Fig. 1.**
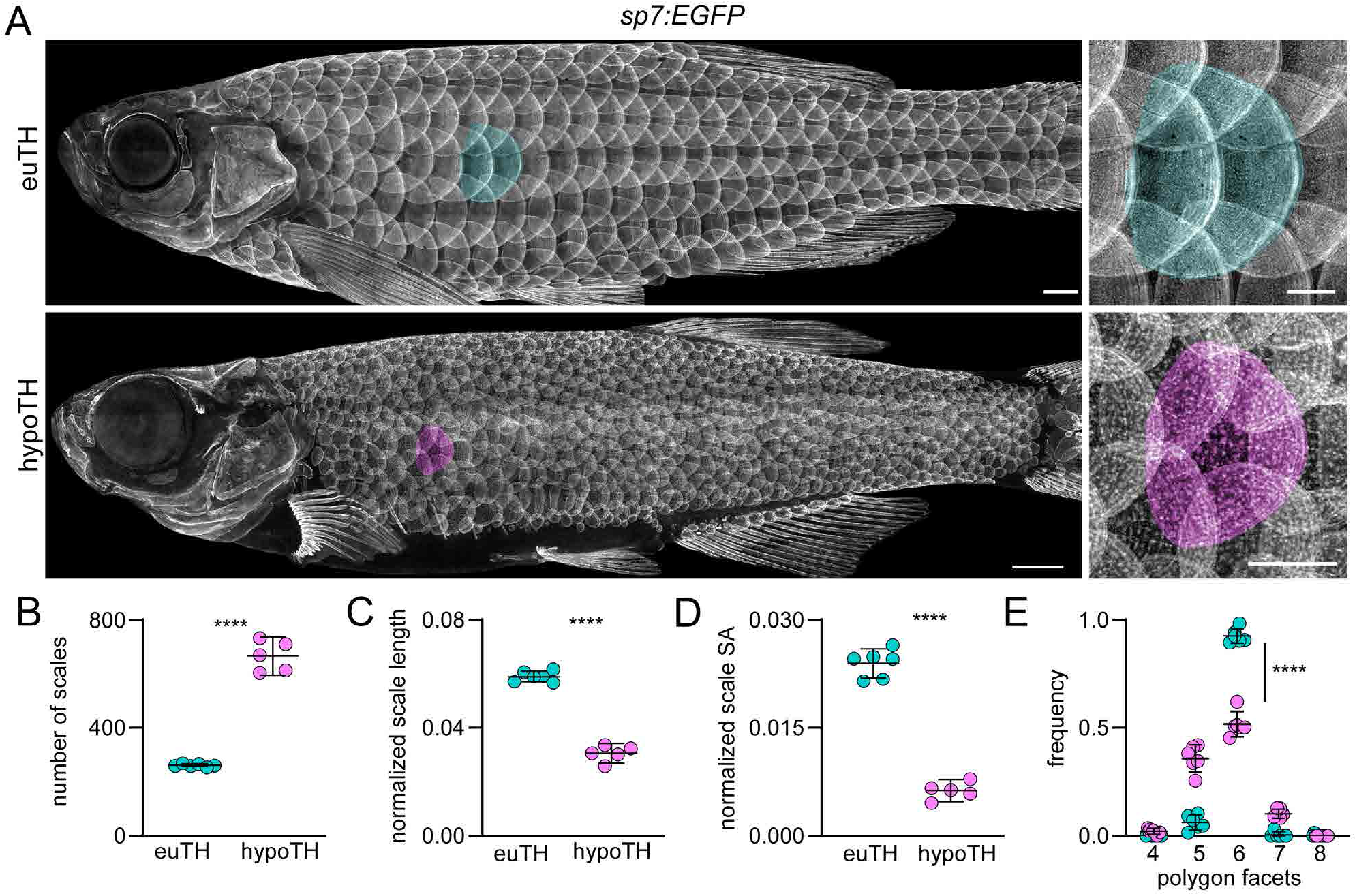
Thyroid hormone is necessary for normal squamation. (**A**) Dermal skeleton of eu-thyroid controls (euTH, *n*=6) and hypothyroid (hypoTH, *n*=5) fish visualized with *sp7:EGFP*. Representative scales false colored in green (euTH) and magenta (hypoTH). (**B**) hypoTH fish developed more numerous scales than euTH controls (*t_9_*=17.55, *p*<0.0001). (**C**) hypoTH scales were shorter relative to fish body length, when scales were measured from their anterior to posterior margins and these lengths expressed as a proportion of the fish standard length (*t_9_*=19.22, *p*<0.0001; *n*=6 euTH, 5 hypoTH). (**D**) hypoTH scales occupied a significantly smaller portion of the skin surface area (SA) compared with euTH controls (*t_9_*=17.58, *p*<0.0001). (**E**) hypoTH squamation was disorderly compared to euTH controls, quantified by Voronoi tessellation analysis of pattern hexagonality. Each bin, 4–8, shows the proportion of scales bounded by that number of immediate neighbors. The frequency of scales bounded by the modal number of 6 neighbors in euTH animals vs other numbers of neighbors differed between euTH and hypoTH individuals (χ^2^=310.72, d.f.=1, *p*<0.0001, *N*=396 scales of 6 euTH individuals and *N*=1262 scales of 5 hypoTH individuals; ****, t_9_=11.64, *p*<0.0001). Proportions were arcsin-transformed for analysis (Sokal and Rohlf, 1981). Bars in plots are means ± 95% CI. Scale bars, (A) 1 mm.

**Fig. 2.**
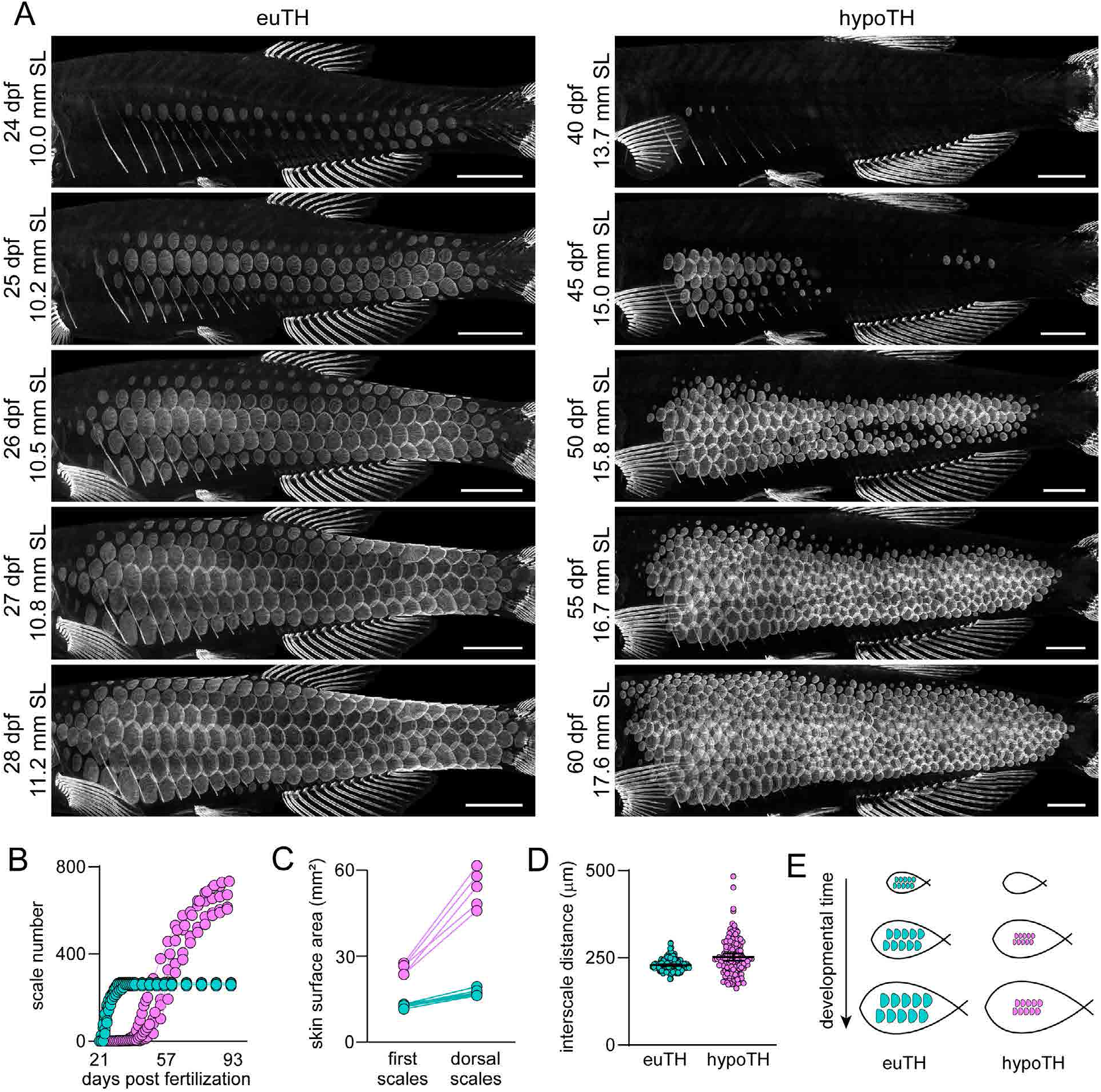
Thyroid hormone regulates squamation onset and progression. (**A**) Timeseries of *sp7:EGFP* expressing scales showing single representative euTH and hypoTH individuals (*n*=6, *n*=5, respectively). In the euTH individual, squamation spread to the dorsal midline in four days, while this process took 20 days in the absence of thyroid hormone. (**B**) The number of scales increased rapidly in euTH fish until reaching the adult complement. In hypoTH fish, scales were added over a long period of time, leading to more total scales than in euTH (*n*=6 euTH, 5 hypoTH). (**C**) Skin surface area expanded dramatically during the slow squamation of hypoTH fish (*n*=6 euTH, 5 hypoTH). (**D**) The distances between anterior–posterior neighbor scales were slightly, but significantly greater in hypoTH fish relative to euTH (*F*_1,13.3_=8.0 *p*=0.014; *n*=10 measurements from each of 6 euTH and 5 hypoTH individuals with individual variation treated as a random effect in mixed model ANOVA; distances were *ln*-transformed for analysis to control for increased variability at larger distances). Bars, means±95% CI. (**E**) Cartoon showing how delayed squamation coupled with a nearly normal interscale distance leads to increased scale density in adults. Scale bars, (A) 1 mm.

**Fig. 3.**
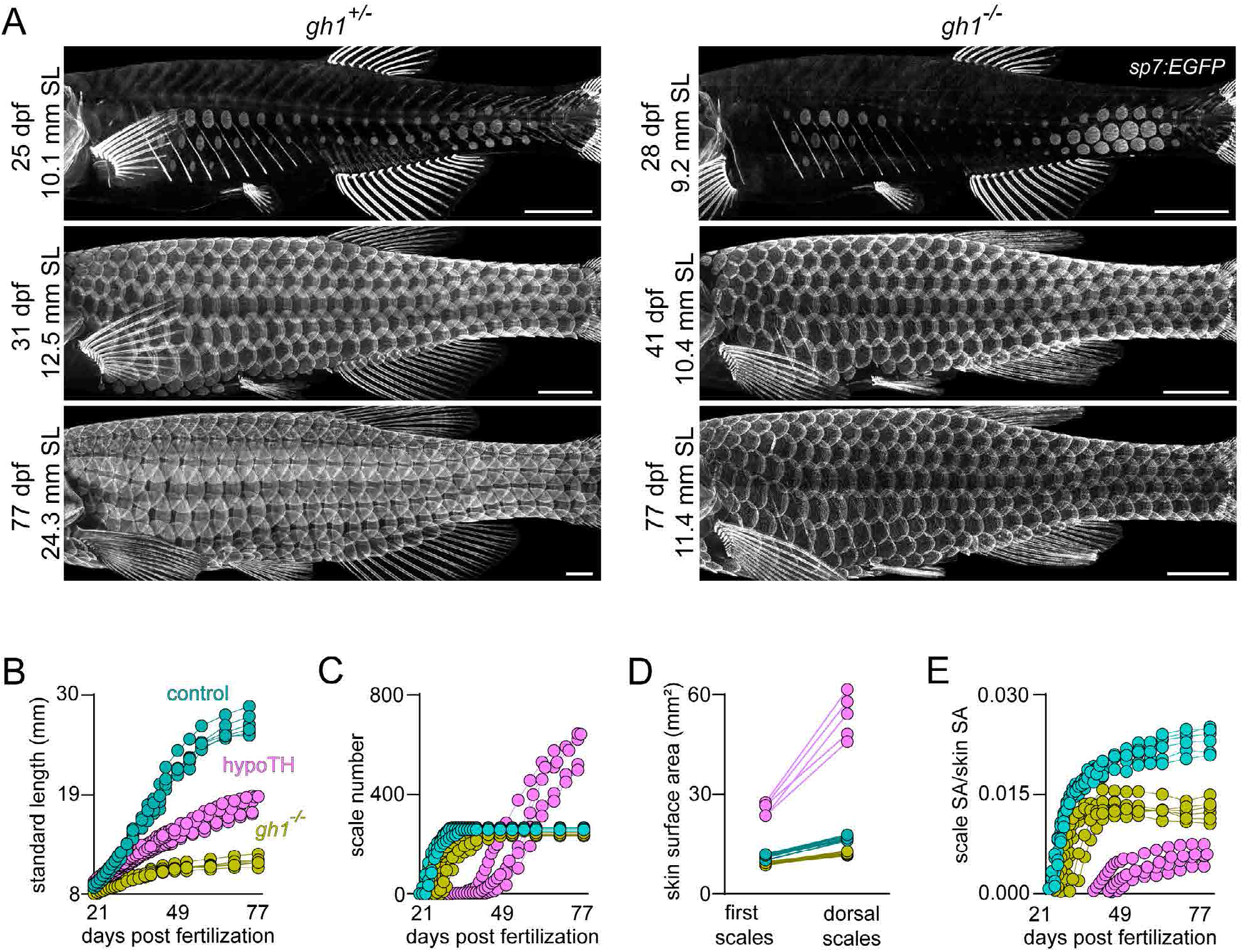
Normal patterning of squamation does not depend on body growth rate. (**A**) Mutants homozygous for a presumptive null allele of *gh1* developed squamation similar to heterozygous siblings, visualized in *sp7:EGFP* transgenics in individual animals over time (*n*=8 each genotype). (**B**) *gh1* mutants grew more slowly than heterozygous siblings or hypoTH fish. (**C**) *gh1* mutants added scales more slowly than heterozygous controls (*n*=8, each condition), but more quickly than hypoTH animals (**D**). *gh1* mutants had less skin surface area expansion than heterozygous siblings (*n*=8), or hypoTH individuals. (**E**) Scale surface area (SA), as a proportion total skin SA over time shows that gh1 mutants have a specific defect in allometric scale growth, despite normal squamation pattern. hypoTH data in C and D is replotted from Fig. 1 for reference. Scale bars, (A) 1 mm.

**Fig. 4.**
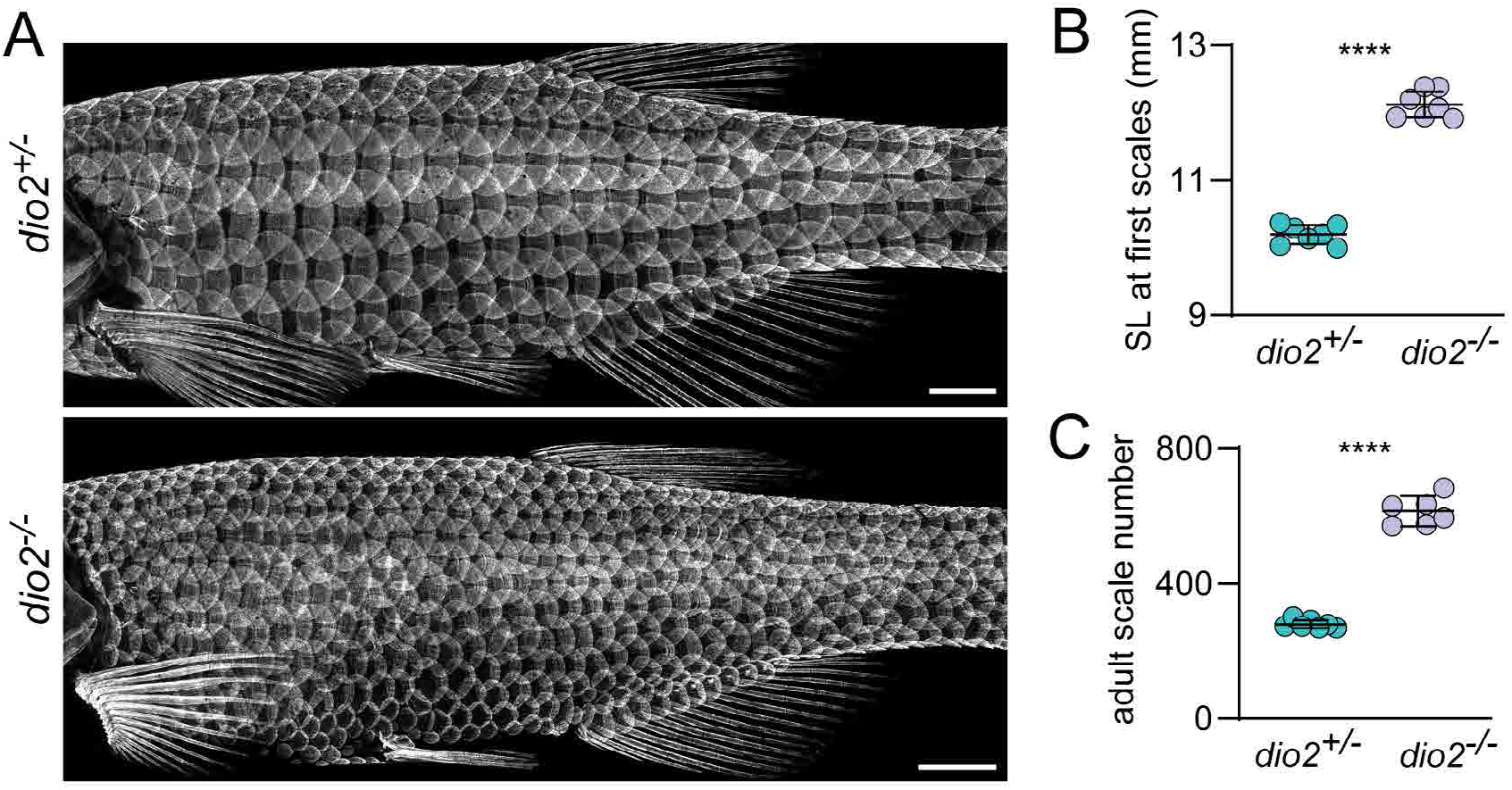
Dio2 is necessary for normal squamation pattern. (**A**) Loss of Dio2 function led to more numerous, smaller scales; similar to fish lacking thyroid glands. Visualized in *sp7:EGFP* transgenics (*n*=7, each genotype). (**B**) Similar to thyroid-ablated *tg:Venus-NTR* hypoTH fish, *dio2* mutants began developing scales later, at a significantly larger standard length (SL; *t*_12_=20.11, *p*<0.0001; *n*=7 each genotype). (**C**) Like thyroid-ablated hypoTH fish, change in timing and size at initiation of scale development was associated with an increased number of adult scales (*t*_12_=19.84, *p*<0.0001). Bars in plots indicate means±95% CI. Scale bars, (A) 1 mm.

**Fig. 5.**
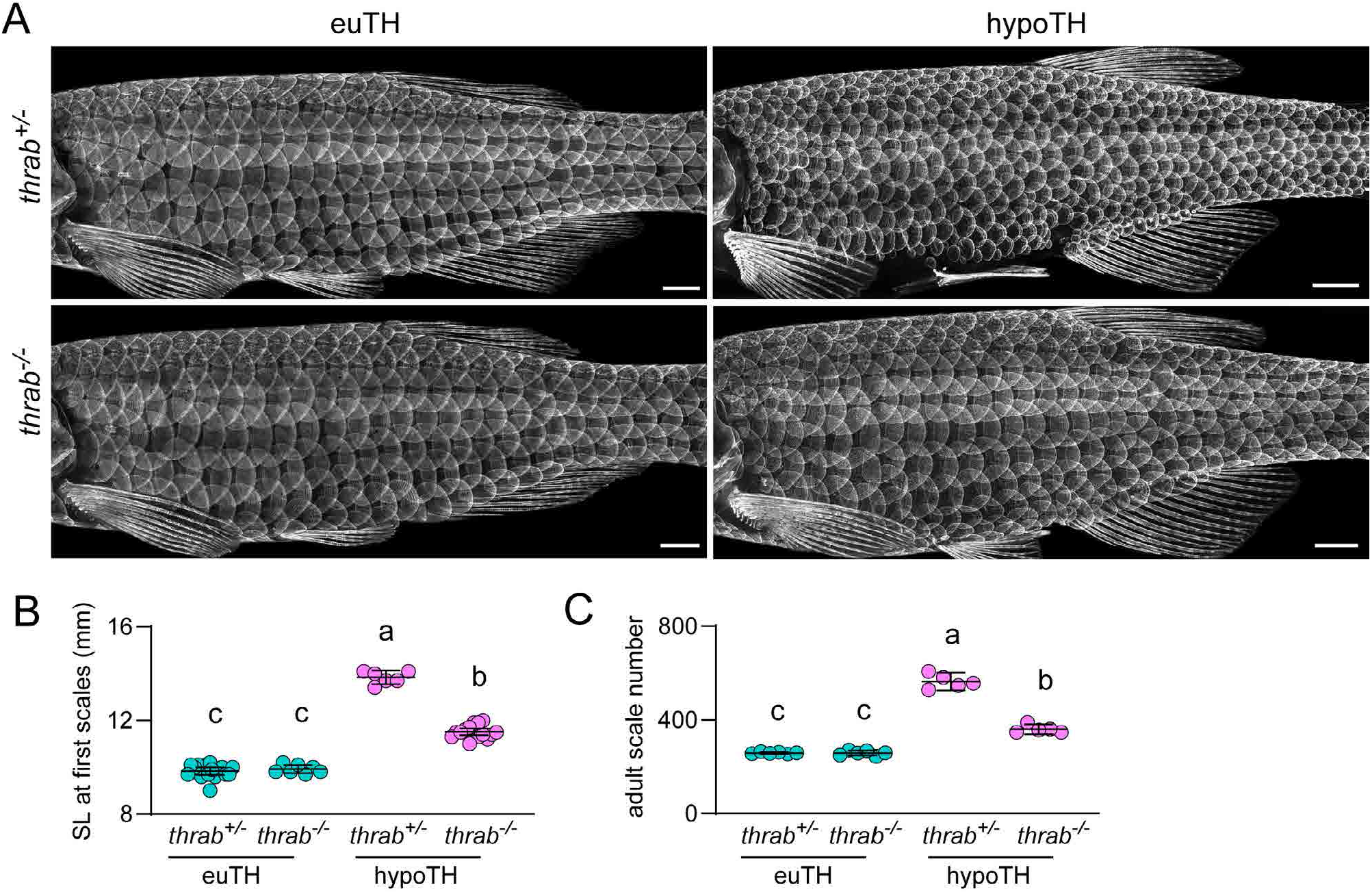
TH regulates squamation by relieving repressive activities of unliganded TRs. (**A**) Euthyroid (euTH) individuals homozygous for a presumptive null allele of thyroid hormone receptor gene *thrab* had an overtly normal pattern of squamation. Whereas hypoTH individuals heterozygous for the *thrab* mutation developed numerous, miniaturized scales, hypoTH animals homozygous mutant for *thrab* had a normal pattern of squamation. Visualized in *sp7:EGFP* transgenics (*n*≥6 each genotype). (**B**) In euTH fish (*n*=16 heterozygotes, *n*=7 homozygous mutants), loss of *thrab* did not significantly affect standard length (SL) at squamation onset whereas in hypoTH fish (*n*=6 heterozygotes, *n*=16 homozygous mutants) loss of *thrab* led to a significant rescue towards the SL at squamation onset of euTH fish. Shared letters above groups indicate means that were not significantly different from one another (*p*>0.05) in *post hoc* Tukey-Kramer comparisons (overall ANOVA, genotype x TH status interaction *F*_1,41_=183.4, *p*<0.0001) (**C**) *thrab* mutation did not affect adult scale number in euTH fish (*n*=6 each genotype), but consistent with the rescue of delayed squamation onset, *thrab* mutation significantly rescued adult scale number in hypoTH fish (*n*=5 each genotype) towards wild-type levels. Scale bar, (A) 1 mm.

**Fig. 6.**
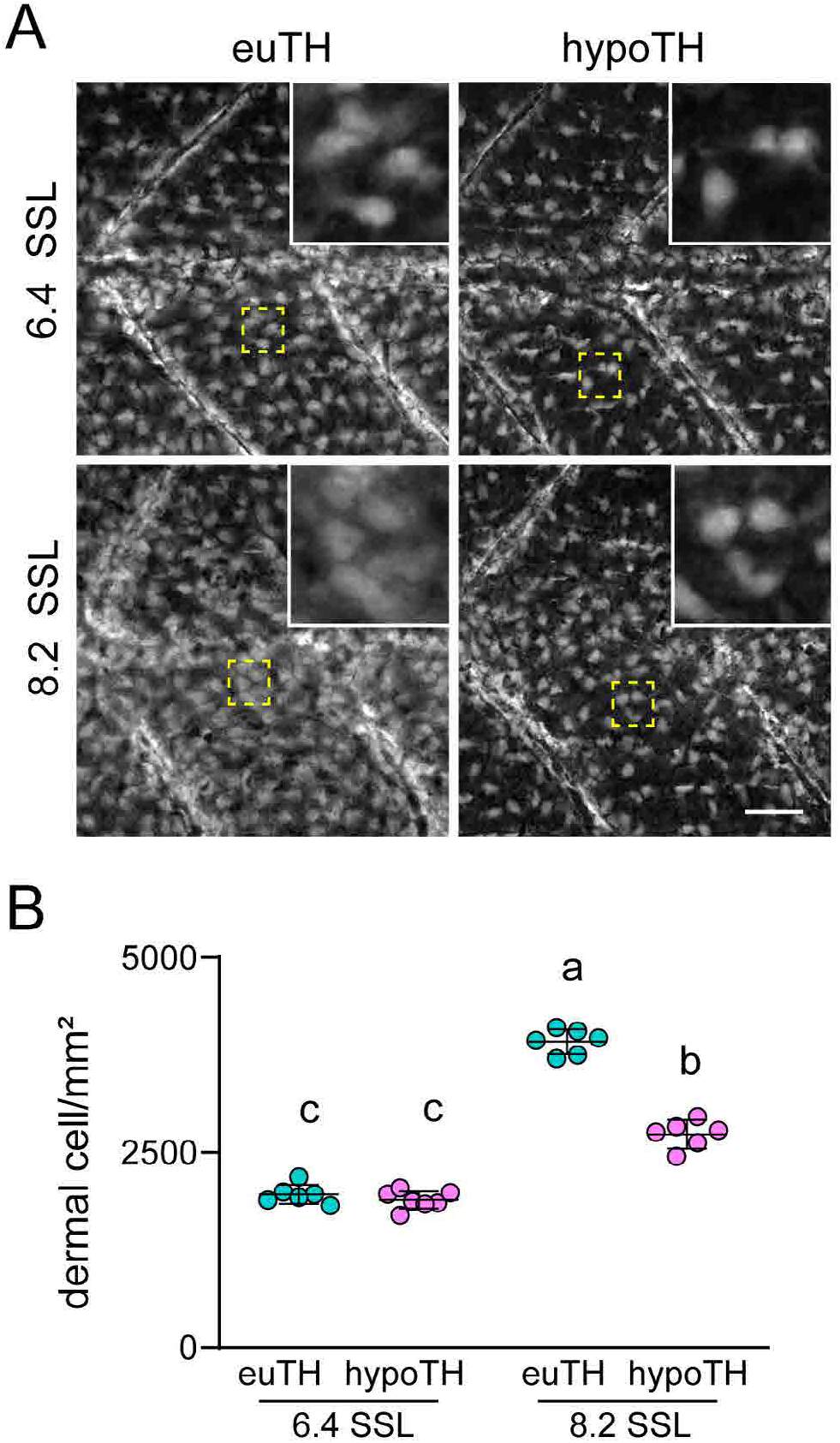
TH regulates densification of the primary hypodermis. (**A**) Primary hypodermal cells, visualized by imaging *ET37:EGFP* transgenic fish, have normal density in hypoTH individuals at 6.4 SSL. By 8.2 SSL, the primary hypodermis of hypoTH individuals appeared sparser than euTH controls. (**B**) Dermal cell density (cell/mm^2^) was similar in euTH and hypoTH fish at 6.4 SSL. By 8.2 mm SSL, euTH fish had significantly denser dermis than hypoTH siblings. Shared letters above groups indicate means not significantly different in *post hoc* Tukey Kramer comparisons (TH condition x stage interaction *F*_1,10.4_=109.1, p<0.0001; mixed model ANOVA with individuals treated as random effect). Scale bar, (A) 100 μm.

**Fig. 7.**
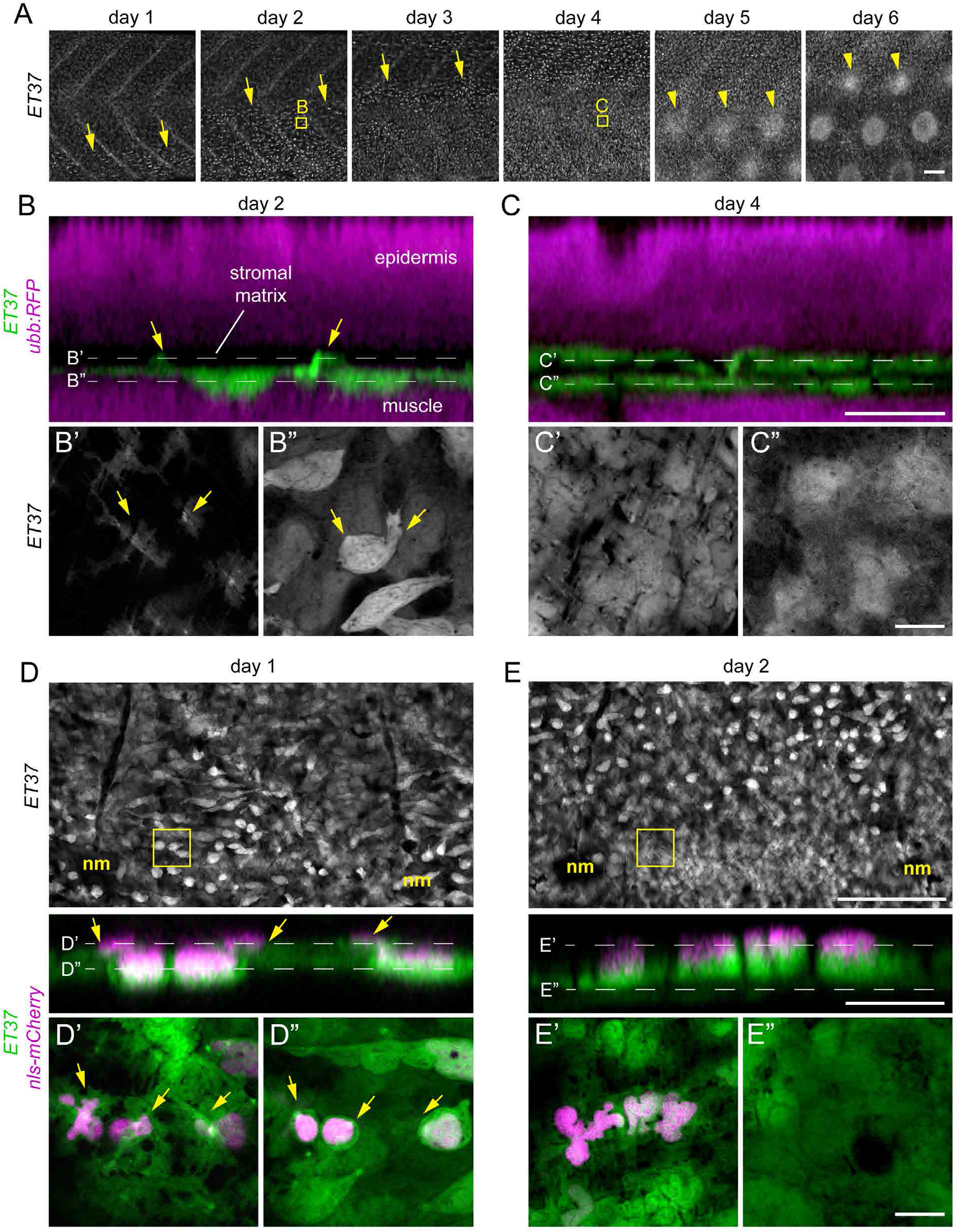
Post-embryonic dermal morphogenesis involves superficial directed migration of primary hypodermal cells. (**A**) Time series of late larval dermal morphogenesis visualized in *ET37:EGFP; ubb:RFP* transgenics. RFP channel omitted for clarity. On day 1 of imaging the first migratory dermal cells appeared near the ventral side of the skin as rounded cells with relatively intense *ET37:EGFP* expression (yellow arrows). Over the following three days, dermal migration swept dorsally and was completed by day 4. Scale papillae (yellow arrowheads) developed after dermal stratification had completed and followed a similar ventral–dorsal wave. Images are from a single representative individual (*n*=8 fish total). (**B**,**C**) Super-resolution imaging of boxed regions in A, showing early (B) and completed (C) dermal stratification. Initially (day 2), the subset of primary hypodermal that express elevated *ET37:EGFP* extended (yellow arrows) toward the epidermis (labeled with *ubb:RFP*). Two days later (day 4) the dermis had been remodeled into a multilayer tissue. (**D**,**E**) Repeated super-resolution imaging of dermal cells expressing a nuclear label (*7xTCF-siam:nls-mCherry*) revealed pre-migratory dermal cells (yellow arrows) that subsequently translocated into a superficial layer within 24 hours. Scale bars, (A) 100 μm, (in C, for B and C), 10 μm.

**Fig. 8.**
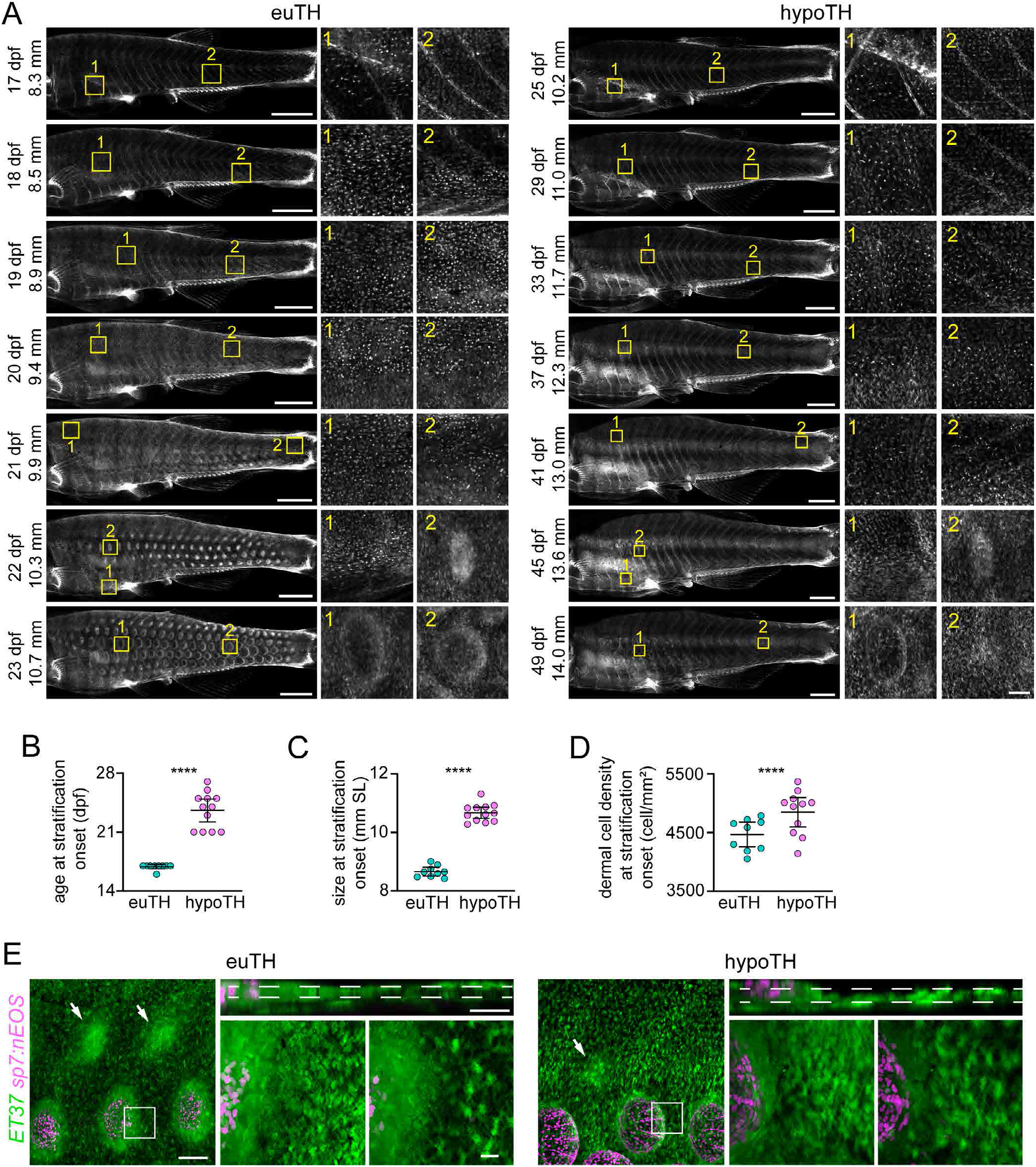
TH regulates dermal migration. (**A**) *In-toto* imaging of dermis in *ET37:EGFP* transgenics revealed a transition between a single-layered primary hypodermis to scaled skin over seven days [17–23 days post fertilization (dpf)]. Beginning at 17 dpf migratory dermal cells were apparent over the ribs (inset 1) but not above the anal fin (inset 2). A similar distribution was observed in hypoTH fish at 25 dpf. Dermal migration occurred in a wave that spread first posteriorly and then dorsally in both euTH and hypoTH fish, followed by scale papilla formation. Although the same qualitative sequence occurred in hypoTH individuals, it progressed more slowly. (**B**) Dermal migration was initiated in significantly older hypoTH fish compared with euTH controls (*t_19_*=9.19, *p*<0.0001; *n*=9 euTH, 12 hypoTH). (**C**) Owing to delayed initiation, skin surface area was significantly larger in hypoTH fish at the onset of dermal migration relative to controls (*t_19_*=18.45, p<0.0001). (**D**) Density of primary hypodermal cells at the onset of dermis migration was more variably and slightly increased in hypoTH fish compared with euTH controls *t_18_*=2.56, *p*=0.02). (**E**) dSFCs differentiated in the superficial dermis in both euTH and hypoTH fish indicated by *sp7:nucEOS* expression (magenta). Bars in B–D, means±95% CI. Scale bars, (A) 500 μm for whole skin overviews or 100 μm for details, (E) 100 μm low magnification and 20 μm for details.

**Fig. 9.**
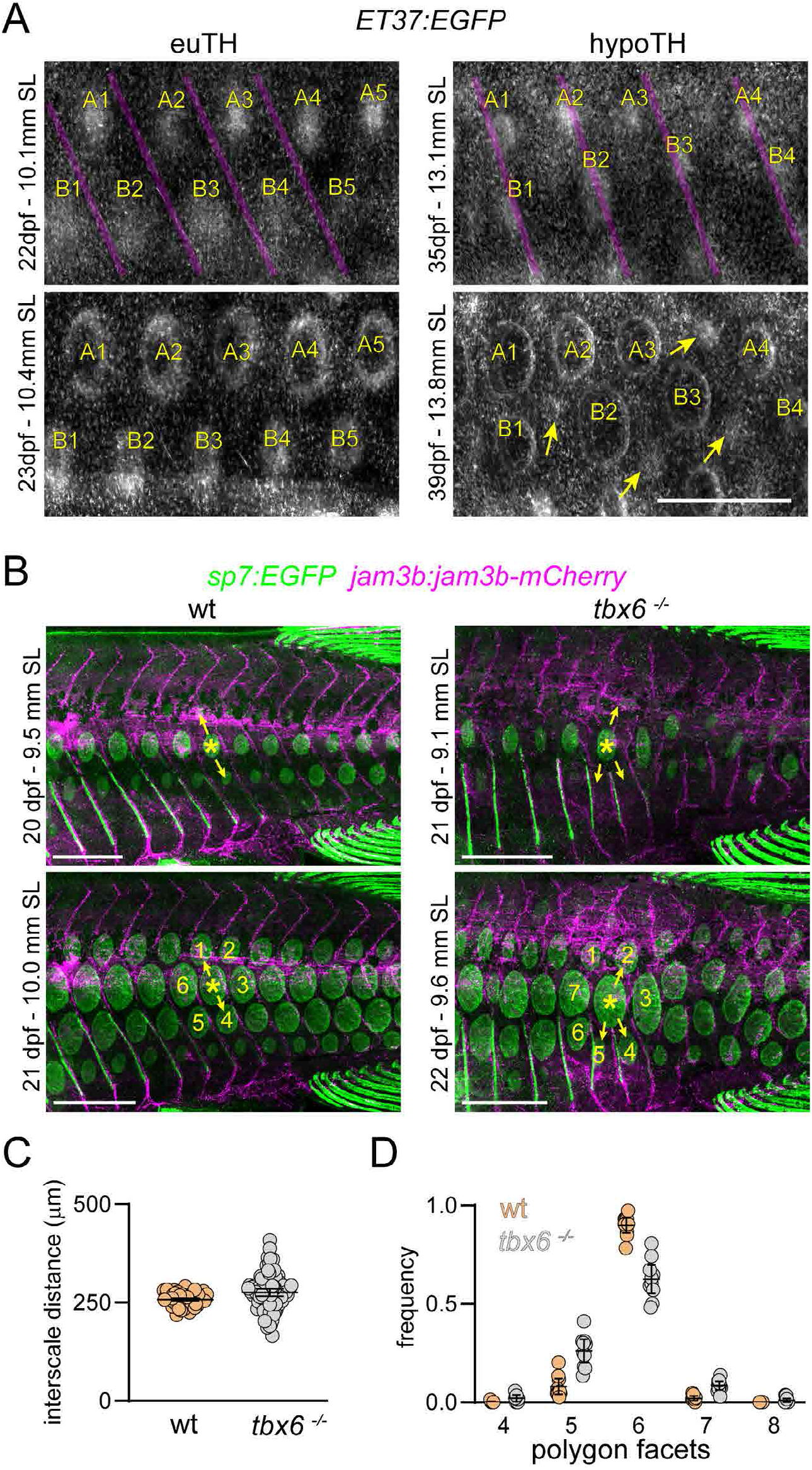
Normal squamation pattern is coupled to segmented myotomes. (**A**) In euTH control fish, scales (numbered by column and lettered by row) formed in the center of myotomes bounded by vertical myosepta (magenta overlay). In hypoTH individuals, initial scale papillae (numbered and lettered) developed closer to myosepta, and multiple scales formed above some myotomes (e.g., A1, A2, A3 over two myotomes). As patterning unfolded, new scale papillae (yellow arrows) appeared within the same rows as initial papillae, associated with increasingly disordered patterns. (**B**) *tbx6* mutants, which mis-patterned vertical myosepta (labelled with *jam3b:jam3b-mCherry*), had disrupted patterns of squamation, visualized by *sp7:EGFP*. In both wild-type control animals and *tbx6* mutants, myosepta predict the location of successive scale papillae (yellow arrows). In the mutant, an apparently branched myotome lead to incusion of an extra papilla which disrupted hexagonal patterning. (**C**) Anterior-posterior distances between early scale papillae were slightly increased and more variable in *tbx6* mutants (effect of TH status, *F*_1,9.3_=6.7, *p*=0.0289; mixed model ANOVA with individual fish treated as a random effect and distances In-transformed for analysis to stabilize variance of residuals among groups; *n*=10 measurements from 5 sibling controls and 8 mutants). (**D**) Voronoi tessellation analysis showed a significant reduction in pattern fidelity in *tbx6* mutants relative to wild-type, as assessed by the incidence of scales bounded by the modal number 6 neighbors found in wild-type vs. numbers of neighbors other than 6 (χ^2^=110.7, d.f.=1, *p*<0.0001, *N*=572 scales of 10 sibling controls and *N*=551 scales of 10 *tbx6* mutants). Bars in plots, means±95% CI. Scale bars, (A) 500 μm, (B) 1 mm.

### 2.6. Image analysis

All measurements were made using FIJI (Schindelin et al., 2012). Surface area measurements were performed on maximum projection images and therefore systematically underestimate the true skin surface area. Scale size measurements were made on representative scales over the ribs (false color in Fig. 1A). 3–4 scales were measured per time point. Voronoi tessellation analysis was performed in FIJI using the Voronoi binarization tool followed my manual scoring of polygon frequency.

### 2.7. Statistics

Statistical analyses were performed using Prism software (Graphpad) or JMP Pro 15 (SAS Institute). In parametric analyses, residuals were inspected for normality and homoscedasticity and variables transformed as needed (described in main text). For repeated sampling within individual fish, we treated individual variation as a random effect in mixed model analyses to avoid pseudoreplication.

## 3. Results

### 3.1. TH regulates scale patterning

Thyroid hormone is necessary and sufficient for the entire suite of morphological changes that characterize amphibian metamorphosis (Brown and Cai, 2007; Buchholz, 2017; Shi, 2000). For elucidating roles of TH in zebrafish, we previously generated a transgenic line, *Tg(tg:Venus-NTR)^wpr8Tg^*, that can be used to efficiently and permanently ablate the thyroid at early larval stages, thereby generating completely hypothyroid (hypoTH) individuals. A preliminary description of hypoTH fish indicated defects in pattern and timing of scale development (McMenamin et al., 2014). To assess this phenotype in more detail, we ablated thyroid glands of fish doubly transgenic for *tg:Venus-NTR* and *sp7:EGFP* (DeLaurier et al., 2010) at 5 days post-ferilization (dpf) and reared hypoTH and euTH control larvae through scale development. *sp7:EGFP* is expressed in cells that deposit calcified matrix, such as skeletal osteoblasts and dermal scale forming cells (dSFCs), allowing scale morphology and patterning to be imaged in live animals (Aman et al., 2018; Cox et al., 2018; Iwasaki et al., 2018; Rasmussen et al., 2018). This analysis showed that adult hypoTH fish develop twice as many scales that were half the size of scales in euTH siblings (**Fig. 1A–D**). Miniaturized scales in hypoTH skin nevertheless retained a seemingly normal cycloid scale morphology.

Zebrafish scales, like other vertebrate skin appendages, are arranged in a well-ordered hexagonal grid. To evaluate the fidelity of scale pattern in hypoTH fish, we employed Voronoi tessellation analysis, which has been used to assess the arrangement of hair follicles in mouse embryos and feather buds in avian embryos (Cheng et al., 2014; Ho et al., 2019). This analysis generates a tessellation map that embeds each pattern element, individual scales in this case, in the largest possible regular polygon. In a perfectly ordered pattern, 100% of pattern elements would be bounded by a hexagon. Consistent with a highly ordered arrangement, 93% of scales in euTH fish had six neighbors. This arrangement was true of only 46% of scales in hypoTH fish, indicating significantly less ordered patterns than euTH controls (**Fig. 1E**). Together, these analyses show that TH is necessary for normal squamation in zebrafish.

### 3.2. TH regulates the timing of squamation onset

TH elicits a bewildering array of responses in different organismal contexts. Depending on the organism, tissue and cell type, TH can promote or inhibit morphogenetic cell behaviors, such as proliferation and migration, that contribute to morphogenesis (McMenamin et al., 2014; Saunders et al., 2019; Shi, 2000; Tata, 1998; Volkov et al., 2020). TH also modulates expression and activity of other signaling pathways, including the Wnt/β-catenin and Fgf pathways, which regulate the patterning of vertebrate skin appendages, including zebrafish scales (Aman et al., 2018; Dalle Nogare and Chitnis, 2017; Harris et al., 2008; Kim and Mohan, 2013; Painter et al., 2012; Rohner et al., 2009; Sire and Akimenko, 2004; Skah et al., 2017; Widelitz et al., 1996). These observations suggest a model in which TH regulates the patterning of scales by affecting local signaling interactions required to set the distance between nascent scale papillae. If this model is true, we predicted that distances between scale papillae should be about half as much in hypoTH fish as euTH fish, given that hypoTH fish have approximately twice the number of scales as euTH fish (**Fig. 1A,B**).

An alternative model of TH activity in this context would emphasize timing: TH regulates the rate of post-embryonic development in other contexts, and could similarly influence the timing of squamation (McMenamin et al., 2014; Saunders et al., 2019; Shi, 2000; Woltmann et al., 2018). If timing rather than distance between papillae depends on TH, we predicted that scales in hypoTH fish should form later than in euTH controls, in skin with approximately twice as much surface area.

To evaluate these predictions, we imaged scale development repeatedly throughout the larva-to-adult transition across entire skins of individual euTH and hypoTH fish expressing *sp7:EGFP*. We found that TH regulated the timing of squamation relative to overall developmental progression and body growth. euTH control fish developed scales beginning at ~21 days post-fertilization (dpf) and had nearly completed this process within four days, similar to previous observations (Aman et al., 2018; Sire and Akimenko, 2004). In hypoTH fish, however, both onset and progression of squamation were delayed: initial scales did not form until ~40 dpf and complete squamation of the ventrum, the final region to develop scales in euTH fish, had yet to be achieved by ~100 dpf (**Fig. 2A,B**; **Movie 1,2**). Due to this delay, squamation was initiated in a much larger skin in hypoTH than in euTH fish (**Fig. 2C**). The anterior–posterior distances between nascent scale papillae were similar in euTH and hypoTH individuals, though these distances were somewhat more variable in hypoTH individuals (**Fig. 2D**). TH therefore promotes an earlier and more abrupt period of squamation than occurs in the absence of this hormone.

These observations suggest that altered scale patterning in hypoTH fish can be largely explained by delayed squamation, combined with the maintenance of nearly normal inter-scale distances: since hypoTH skin was larger at squamation onset but distances between scales were of similar magnitude, more scales were required to cover hypoTH skin than euTH skin (**Fig. 2E**). The phenotype of increased scales per unit area in hypoTH fish is therefore most likely a secondary consequence of developmental delay in their formation.

This live imaging experiment serendipitously revealed heterogeneities in TH-responsiveness for additional traits as well. During the zebrafish larva-to-adult transition, the larval fin fold is resorbed and the pelvic fins show rapid allometric growth (Parichy et al., 2009). Resorption of the larval fin fold was delayed in hypoTH individuals, demonstrating for the first time in zebrafish a role for TH in destruction of a larval trait (**Fig. S1A–C**). In contrast to formation of scales and destruction of the fin-fold, the pelvic fin grew more slowly in hypoTH fish, yet retained its normal allometry relative to the body (**Fig. S1A,D,E; Movies S1,S2**). The eye grew at the same rate in hypoTH and euTH fish, demonstrating TH-independence of its growth (**Fig. S2; Movies S1,S2**).

Together these analyses of ontogenetic changes in individual fish indicate that different traits are differently regulated by TH. Whereas the eye grew normally in the absence of TH, growth of the pelvic fin, disappearance of the fin fold, and initiation and progression of squamation were each delayed in hypoTH individuals. Of these alterations, only delayed squamation led to an overt patterning defect in the adult.

### 3.3. Scale patterning depends on developmental coupling rather than absolute growth rate

In amphibians, TH is essential for morphogenetic changes at metamorphosis—limb development, intestinal remodeling, tail resorption and other alterations—yet TH is not required for growth of the body itself, which is thought to be regulated independently by growth hormones (Brown and Cai, 2007; Buchholz, 2017; Shi, 2000; Tata, 1998). In zebrafish, live imaging here, and observations of prior studies, indicate that TH promotes the development of some specific traits, like scales, but is also necessary for somatic growth overall, as hypoTH fish increase in size more slowly than euTH fish (Hu et al., 2019; McMenamin et al., 2014). Live imaging of squamation in euTH and hypoTH fish strongly suggests that TH regulates scale patterning by coupling scale papilla formation to body growth (**Fig. 2**). Therefore, we predicted that fish compromised in somatic growth for reasons other than TH insufficiency could well exhibit squamation timing and pattern equivalent to wild-type. Accordingly we examined *growth hormone 1 (gh1)* mutants, which have markedly reduced growth rates during the larva-to-adult transition, leading to relatively diminutive adults (**Fig. 3A,B; Fig. S3**) (McMenamin et al., 2013).

We repeatedly imaged *sp7:EGFP* expression in individual *gh1* mutants and wild-type siblings throughout squamation and found scale patterning of *gh1* mutants to be grossly normal (**Fig. 3A**), despite these fish having growth rates even slower than hypoTH fish (**Fig. 3B**). Further analyses showed that *gh1* mutants had a slight delay in absolute timing of both onset and completion of squamation (**Fig. 3C**) but that slow squamation remained coupled to the slowly expanding skin (**Fig. 3D**), ultimately leading to a squamation pattern that resembled the wild type (**Fig. 3A**). This condition contrasts with squamation in hypoTH individuals, in which body growth and squamation timing were decoupled, leading to development of scales in much larger skin (**Fig. 2A-C**).

Individual scales initially grow allometrically followed by isometric growth as they overlap and the animals approach adulthood (Goodrich and Nichols, 1933; Sire and Akimenko, 2004). Our live imaging experiment showed that although scale patterning was not affected by *gh1* loss, the mutants developed proportionally smaller scales following an early and abrupt shift to isometric growth, consistent with a specific role for GH1 in promoting growth of bony tissues (**Fig. 3E)** (Slootweg, 1993). These results support a model in which normal squamation pattern requires coupling between scale papilla formation and body growth, rather than absolute timing of growth or squamation. We next turned our attention to the molecular and cellular mechanisms underlying TH dependent coupling of body growth and skin morphogenesis.

### 3.4. Scale development is regulated by TH nuclear hormone pathway in skin

TH can regulate gene expression in target cells via genomic or non-genomic signal transduction pathways (Davis and Davis, 1996; Hammes and Davis, 2015; Shi, 2000; Tata, 1998). In genomic TH signaling, the circulating TH prohormone T4 (thyroxine) is converted to the more biologically active T3 (triiodothyronine) in target cells by a specific deiodinase enzyme, Deiodinase 2, encoded by *dio2*. Following deiodination, T3 interacts with TH nuclear receptors (TRs), leading to conformational changes that strongly decrease TR affinity for transcriptional co-repressors and increase affinity for co-activators to regulate gene expression (Bianco and Kim, 2006; Brent, 2012; Buchholz et al., 2003; Han et al., 2020; Hörlein et al., 1995). By contrast, the most well characterized non-genomic TH pathway involves high affinity binding of T4 to specific cell surface integrin receptors (Bergh et al., 2005). If scale development is regulated via genomic TH signaling, we predicted that loss of Dio2 activity would lead to a scale patterning phenotype similar to hypoTH fish. Conversely, if scale development depends primarily on T4 mediated non-genomic signaling, loss of Dio2 may not effect squamation. To discriminate among these possibilities, we generated a *dio2* mutant (**Fig. S4)** and assayed scale development by imaging *sp7:EGFP*. This analysis revealed that, indeed, loss of *dio2* causes later-developing and more numerous scales, similar to hypoTH fish (**Fig. 4**).

To additionally test whether TRs regulate squamation, we assayed *sp7:EGFP* expression throughout scale development in mutants for *thyroid hormone receptor ab (thrab)* that were either euTH (intact thyroid) or hypoTH (thyroid ablated) (Saunders et al., 2019). These analyses showed that loss of *thrab* in euTH fish did not result in delayed squamation or scale patterning defects, however, this mutation was sufficient to rescue both the timing of squamation and scale pattern in hypoTH fish (**Fig. 5**). This result suggests that unliganded TRs repress the onset and progression of scale development in control animals, and that this repression is relieved in the presence of T3. In the *thrab* mutant, target loci are derepressed owing to absence of the receptor, despite lack of T3. Together with analyses of *dio2* mutants, these results suggest genomic transduction is the likely molecular mechanism underlying TH signaling in scale development.

TH circulates widely and TH signaling could potentially regulate squamation directly, by signaling to skin cells, or indirectly by regulating gene expression in another organ system such as the cutaneous vasculature or underlying muscles. To evaluate whether skin itself is likely to be a direct target of TH signaling we examined the expression of TH transporters, receptors, and co-receptors by *in-situ* hybridization. These analyses showed that transcripts for all three classes of factors were present in skin including scale primordia, indicating that these cells are likely competent to respond directly to TH (**Fig. S5**). To test this possibility, we assayed skin expression of a synthetic TH reporter transgenic line, *TRE:EGFP*, in which tandem consensus TH Response Elements (TRE) drive EGFP expression (Matsuda, 2018). These analyses confirmed TH-dependent reporter expression in both epidermis and dermis (**Fig. S6**), supporting a model in which the skin itself is a direct target of TH.

These analyses show that squamation is largely regulated by genomic TH signaling and such signaling is likely to take place in the skin itself, where all major classes of TH pathway genes were expressed and TH signaling activity was detectable with a synthetic reporter transgenic. Though our findings cannot exclude additional roles for non-genomic TH signaling or indirect effects of TH via other tissues, these results strongly suggest that skin is a direct target of genomically mediated TH signaling in zebrafish.

### 3.5. TH regulates early dermal development

Our analysis suggested that scale development depends directly on TH, however, measurements of TH (T3 and T4) over the course of post-embryonic development have shown that TH levels increase much earlier in development, with a peak occurring at 6.4 standardized standard length (SSL) (Hu et al., 2019), roughly one week prior to scale development, which begins at 9.6 SSL under standard conditions (Aman et al., 2018; Parichy et al., 2009). Since normal scale development is preceded by extensive dermal remodeling (Le Guellec et al., 2004; Sire and Akimenko, 2004) we hypothesized that TH is required for earlier events in dermal development that are, in turn, essential for squamation to proceed. Because tests of this hypothesis would be enabled by visualizing dermal cells *in vivo*, we asked whether an enhancer trap reporter line, *ET37:EGFP*, used to visualize dermal cells and fin fold mesenchyme in early larvae (Morris et al., 2018; Parinov et al., 2004; Zhang et al., 2010), might be useful during later larval development as well. To assess whether *ET37:EGFP* is expressed in post-embryonic dermal cells, including dermal cells that contribute to scale development, we sectioned transgenic fish and counterstained with a membrane dye to label all cells. This revealed that *ET37:EGFP* is expressed in all dermal cell subtypes known from histological studies, including the hypodermis (occasionally referred to as the dermal endothelium), stromal cells or fibroblasts, superficial dermal cells, scale papillae, and cells within the maturing scale, though expression was weaker in dSFCs, which express osteoblast markers such as *sp7* and deposit calcified scale matrix itself (**Fig. S7**) (Aman et al., 2018; Iwasaki et al., 2018; Le Guellec et al., 2004; Sire and Akimenko, 2004). Given that dermal cells were marked by *ET37:EGFP* at late stages of larval development, we used this line in combination with thyroid ablation line *tg:Venus-NTR* to assay TH requirements for dermal development.

If TH is required for dermal development, we predicted that a phenotype should be evident in hypoTH fish as early as 6.4 SSL, coinciding with peak TH accumulation (Hu et al., 2019). Imaging *ET37:EGFP* at this stage, we found that both euTH and hypoTH fish had a single layer of dermal cells, which we will call the primary hypodermis to distinguish from the mature hypodermis. At this stage, the dermis constituted a monolayer of primary hypodermal cells in both euTH and hypoTH fish. The density of these cells, measured as the number of cells per unit area, was similar although cells in hypoTH appeared to be individually less uniform in *ET37:EGFP* expression (**Fig. 6A** *upper* and **Fig. 6B**). By 8.2 SSL, the euTH primary hypodermis contained significantly more cells per unit area than hypoTH (**Fig. 6A** *lower* and **Fig. 6B**). This slower accumulation of primary hypodermal cells between 6.4 SSL and 8.2 SSL was the earliest phenotype we could reliably quantify in hypoTH fish, and its timing was concordant with that of the early peak described for TH abundance and presumed availability.

Prior to scale formation, the dermis remodels from a simple monolayer to a stratified tissue comprising a deep hypodermis, stromal fibroblasts, and superficial dermal cells that ultimately differentiate into dSFCs (Aman and Parichy, 2020; Le Guellec et al., 2004; Sire and Akimenko, 2004). Although there is a comprehensive report on the histology and ultrastructure associated with this transformation, cellular dynamics remain incompletely characterized (Le Guellec et al., 2004). To elucidate such dynamics and their potential significance for later squamation, we repeatedly imaged individual euTH fish expressing *ET37:EGFP* to visualize dermal cells and ubiquitously expressed RFP (*ubb:RFP*) (Mosimann et al., 2011; Pan et al., 2013) for histological context throughout the period spanning from primary hypodermal monolayer to scaled skin. Conventional confocal imaging of these individuals revealed that dermal stratification began at 8.2 SSL and proceeded in a ventral-to-dorsal wave over the course of three days, followed closely by a wave of scale papillae formation (**Fig 7A**). To visualize cellular dynamics underlying dermal stratification, we imaged a representative location of these fish over time at superresolution (yellow boxes in **Fig. 7A**). These analyses revealed that a subset of primary hypodermal cells individually increase *ET37:EGFP* expression and extend processes into the collagenous stroma towards the epidermis (**Fig. 7B**) followed closely by appearance of a stratified dermis comprising 2–3 layers of *ET37*:EGFP+ cells (**Fig. 7C**). A subset of primary hypodermal cells with processes extended towards the epidermis were likewise evident in histological sections at 8.6 SSL (**Fig. S7A,B**).

These observations suggested the hypothesis that *ET37:EGFP*+ primary hypodermal cells migrate from basally into the stroma to form more superficial dermal layers. To test this hypothesis, we imaged dermal *ET37:EGFP*+ cells in individual fish additionally transgenic for *7xTCF-siam:mCherry*, which labels a subset of dermal cells with nuclear-localizing mCherry, the sparsity of which allows for re-imaging of the same cells over time. This assay revealed that primary hypodermal cells expressing high levels of *ET37:EGFP* translocated to a more superficial position over the course of a day (**Fig. 7D,E**). These results show that a subpopulation of primary hypodermal cells migrates toward the epidermis to generate a stratified dermis. Remarkably, the dermis transformed from a simple monolayer to a stratified tissue complete with scales in just five days, representing an abrupt morphogenesis of the skin.

Because dermal stratification always preceded scale development (**Fig. 7A,B**), we hypothesized that dermal stratification is regulated by TH, and that a delay in such stratification accounts for the delay in scale formation in hypoTH fish. An alternative possibility would be that dermal stratification occurs normally in hypoTH fish, which in turn would suggest that delayed scale development in hypoTH fish might represent a more specific requirement of developing scale papillae for TH. To discriminate between these possibilities, we exploited the upregulation of *ET37:EGFP* in primary hypodermal cells prior to stratification to assay spatial and temporal patterns of stratification using *in-toto* dermal imaging of euTH and hypoTH fish, beginning just prior to the onset of stratification in euTH fish (7.2 SSL). In addition to *in-toto* dermal imaging, these individuals were imaged at super-resolution, which confirmed that upregulation of *ET37:EGFP* in a subset of hypodermal cells always preceded dermal stratification in both euTH and hypoTH fish (**Fig. S8**). By assaying the entire dermis, rather than a representative area (as in **Fig. 7A**), we found that that the dermal stratification wave—marked by punctate cells expressing high levels of *ET37:EGFP*—begins anterior, above the ribs, and spreads posterior to the caudal fin before spreading dorsally (**Fig. 8A**). The last regions of skin to stratify were just posterior to the head and at the caudal peduncle (euTH, 21 dpf boxes 1 and 2), and along the ventrum (euTH, 22 dpf box 1). This sequence exactly presaged the order of scale appearance (compare euTH 18 dpf and 22 dpf **Fig. 8A**). Dermal stratification followed a similar progression in skin of hypoTH fish, though this process began later and proceeded more slowly, as compared to the skin of euTH fish (**Fig. 8A–C**).

Because primary hypodermal cells increased their numbers more slowly in hypoTH individuals (**Fig. 6**), we hypothesized that migration might be triggered by a threshold number of cells per unit area within the primary hypodermal monolayer. If so, we predicted that although dermal stratification is delayed in hypoTH individuals compared with euTH individuals, the number of primary hypodermal cells per unit area would be similar at the onset of stratification. To test this, we quantified cell number per unit area at the first timepoint featuring *ET37:EGFP* upregulation by primary hypodermal cells. This analysis showed that, indeed, primary hypodermal cell numbers in hypoTH were similar, and in many cases cases greater, than cell numbers in euTH primary hypodermis at migration onset (**Fig. 8D**). To confirm that superficial *ET37:EGFP* cells differentiated into dermal scale-forming cells (dSFCs), we imaged euTH and hypoTH fish expressing *ET37:EGFP* and *sp7:nucEOS*, which labels nuclei of dSFCs. This live imaging experiment showed that, consistent with previous reports that relied on histology and ultrastructure, dSFCs differentiate within papillae that form in the superficial dermis (**Fig. 8E**) (Le Guellec et al., 2004; Sire and Akimenko, 2004).

Together, these analyses show that TH promotes abrupt dermal remodeling, wherein a simple monolayer is converted into a stratified tissue complete with scales in just a few days, and suggest that a delay in the accumulation of superficial dermal cells is likely to be a proximate cause of delayed squamation in hypoTH fish.

### 3.6. TH couples scales to musculoskeletal system

Though a delay in dermis development leading to delayed initiation of squamation is likely to explain the increased density and smaller scales of hypoTH fish, such changes do not obviously explain the disordered arrangement of scales in these fish (**Fig. 1A,E**). Our *in-toto* dermal imaging enabled visualization of myosepta because ET*37:EGFP*+ dermal cells appear to follow myotome contours (**Figs. 6A, 7A, 8A**). These data revealed a striking correspondence between the pattern of scales and the pattern of underlying myotome segmentation in euTH fish, but not in hypoTH fish (**Figs. 8A, 9A**). In euTH fish, scale papillae formed in the center of myotomes, between vertical mysosepta. In hypoTH fish, papillae tended to form towards the myosepta, and two scales occasionally formed within the boundaries of single myotomes. Additionally, hypoTH fish developed intercalary scales between early formed scales, a phenomenon never observed in euTH control fish (**Fig. 9A**, *lower right*). As scales, like other skin appendages, are thought to be regulated by local self-organizing signaling interactions, delayed squamation relative to body growth in hypoTH fish, combined with an interpapillary distance similar to that of euTH fish, may result in larger patterning fields between vertical myosepta that can accommodate multiple scale papillae.

These observations led us to hypothesize that the segmented pattern of myotomes influences the regular pattern of scales during normal development. To test this hypothesis, we generated fish with myoseptal boundary defects by mutating *tbx6*, which is essential for vertical and horizontal myoseptum development in the embryo and early larva (**Fig. S9**) (Nikaido et al., 2002; van Eeden et al., 1996; Windner et al., 2012). We found that *tbx6* mutants lacked myotome segmentation initially, but some myosepta had formed by later larval stages, though vertical myosepta were disorganized compared to wild-type siblings (**Fig. S10**). To test if myotome pattern influences squamation pattern, we examined the distribution of scales in *tbx6* mutants transgenic for *sp7:EGFP* to visualize scales and *jam3b:jam3b-mCherry*, which labels intersegmental vasculature and other myosepta associated cell types (Eom et al., 2021). These analyses confirmed a close correspondence between myotomes and hexagonally patterned scales in wild-type control animals and a markedly less ordered pattern of squamation in *tbx6* mutants, despite these fish having relatively normal scale spacing (**Fig. 9B–D**). Tbx6 regulates marker genes associated with dermomyotome fate in amniotes (Windner et al., 2012), raising the possibility that scale pattern defects in *tbx6* mutants reflected alterations in dermis development, rather than an influence of mis-patterning in the underlying myotomes. To evaluate this possibility, we imaged the dermis *in-toto* throughout the period of dermal stratification which failed to reveal overt defects in initiation or progression of dermal stratification in *tbx6* mutants (**Fig. S11**). By contrast, specific anomalies of scale pattern were often associated with corresponding anomalies in myotome pattern (**Fig. 9B**).

These results show that regularly spaced myotomes constitute a likely pre-pattern that affects patterning of squamation. Together, these analyses suggest that TH, by driving timely scale formation, ensures that squamation is coupled to the segmented musculoskeletal system.

## 4. Discussion

Many animals adopt their adult form over a period of post-embryonic development that can involve extensive morphogenetic remodeling. This can range in degree from the catastrophic metamorphosis of some protostomes and non-bilaterians, to much more subtle alterations in limb proportion in mammals (Kulin and Müller, 1996; McMenamin and Parichy, 2013; Temereva and Tsitrin, 2013). In zebrafish, we have found evidence for extensive TH dependent post-embryonic dermal remodeling. TH stimulated every investigated aspect of post-embryonic dermal development, including accumulation of cells in the primary hypodermis (**Fig. 6**) and later dermal stratification and scale development (**Fig. 8, Fig. S7**).

TH has been shown to regulate the timing, density and size of scales in zebrafish and other species (Bolotovskiy et al., 2018; Levin, 2010; McMenamin et al., 2014; Smirnov et al., 2006). Our work supports and extends these findings, adding mechanistic insights into the mode of TH action and the cellular dynamics that are regulated. We found that TH likely acts directly on skin, although we are unable to rule out non-autonomous roles of other tissues such as endothelial cells, the digestive system or the nervous system. By imaging dermal development at multiple spatial scales, we found that post-embryonic dermal remodeling occurs in a spatio-temporally invariant wave that sweeps across the skin and provides progenitor cells for dSFC differentiation and scale development. We have found that the rate of these events is regulated by TH. The transcriptional targets of TH and mechanisms linking target genes to morphogenetic cell behaviors, such as cell migration and proliferation, remain unknown.

Our work focused on dermal cells, as these are the cells that ultimately deposit bony scale matrix, however, we have not addressed the role of epidermis on TH-dependent dermal morphogenesis. It is possible that some or all of the dermal phenotypes presented here are due to defects in epidermal development that secondarily affect dermal morphogenesis. Dissecting these possibilities will require new tools for tissue-specific manipulation of TH sensitivity.

We additionally show that TH, by regulating the timing of scale development, is necessary to couple individual scales to the underlying segmented musculoskeletal system. In euTH fish, most scales form on top of individual myotomes leading to a correspondence between the squamation pattern and the segmented musculature. In hypoTH fish, delayed scale formation results in development of multiple scales per segment. Scale patterning, like patterning of other vertebrate skin appendages, is thought to rely on local, self-organized interactions involving relatively short range chemical and physical cues (Cooper et al., 2018; Dalle Nogare and Chitnis, 2017; Green and Sharpe, 2015; Hiscock and Megason, 2015; Ouyang and Swinney, 1991). Our results suggest that in euTH fish, the local patterning mechanism interacts with the underlying segmented musculature to produce the adult scale pattern. Due to delayed squamation relative to body growth in hypoTH fish, myotome segments are large enough to accommodate multiple scales with correct spacing. It is tempting to speculate that this arrangement might be optimal for fish armoring, mobility or both.

The density of skin appendages varies widely among diverse vertebrates, from the luxuriously dense fur of a sea lion to the pathetically sparse pelage of a typical human. Intriguingly, it has recently been reported that the timing of feather bud appearance correlates with the density of plumage in adult birds, with feather buds of emu and ostrich embryos appearing later in development than in duck or chicken embryos (Ho et al., 2019). We can imagine that alterations in TH secretion or tissue-sensitivity might contribute to these and similar evolutionary changes.

Beyond informing our understanding of endocrine regulation during post-embryonic skin development, our work has potential implications for environmental toxicology. Ever increasing discharge of endocrine disrupting pollutants is a pressing concern that affects natural environments and human health (Kumar et al., 2020). Although analytical chemistry methods are very sensitive in detecting endocrine disrupting compounds in environmental samples, these methods require advanced instrumentation and sophisticated sample preparation. Furthermore, these methods can only evaluate the presence of known pollutants (Sosa-Ferrera et al., 2013). Quantifying and comparing the density and orderliness of squamation patterns of wild fish collected from various watersheds, or laboratory zebrafish reared in environmental water samples, might provide a low cost alternative for evaluating the impact of thyroid disrupting compounds in the environment. For large species with pigmented scales, this assay would involve simple macro-photography. For smaller species or species that lack scale pigmentation, staining with calcium binding dyes such as Calcein or Alizarin Red would represent convenient and inexpensive alternatives (Iglesias and Rodríguez-Ojea, 1997).

## Acknowledgements

For assistance we thank Marianne Cole and Amber Schwindling for fish care, and Alice and Clara Aman for fastidious supervision of analysis and manuscript preparation. Supported by NIH R35 GM122471 to DMP.

---

**Fig. S1.**
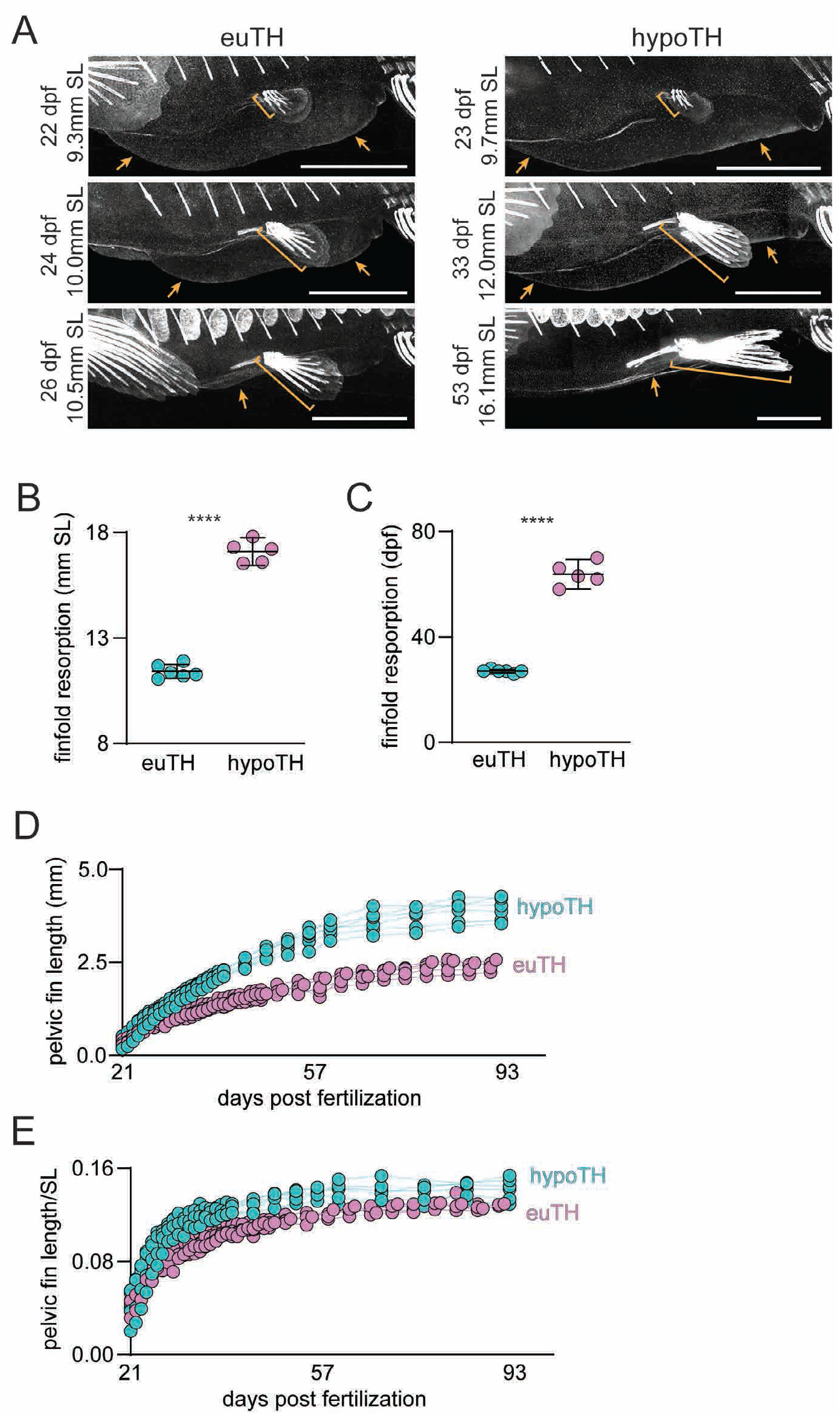
Finfold and pelvic fin morphometrics. (**A**) Representative sequential images from *sp7:EGFP* timeseries showing disappearance of the larval fin-fold (orange arrows) in representative euTH and hypoTH individuals. Though fin fold resorption appeared similar morphologically between euTH and hypoTH, it was markedly delayed in hypoTH fish relative to age, SL [a proxy for stage (Parichy et al., 2009)], and pelvic fin outgrowth (orange bracket). Images are scaled to show similar anatomical region and highlight allometry in the growing fish. (**B, C**) Finfold resorption was delayed in hypoTH animals relative to euTH and completed in significantly larger and older individuals (*t*_9_=20.04, *t*_9_=21.98, both *p*<0.0001; *n*=6 euTH; *n*=5 hypoTH). (**D**) Pelvic fins grew more rapidly in euTH fish compared with hypoTH siblings. (**E**) Following a burst of allometric growth, the pelvic fins grew isometrically in both euTH and hypoTH individuals. Scale bars, (A) 1 mm.

**Fig. S2.**
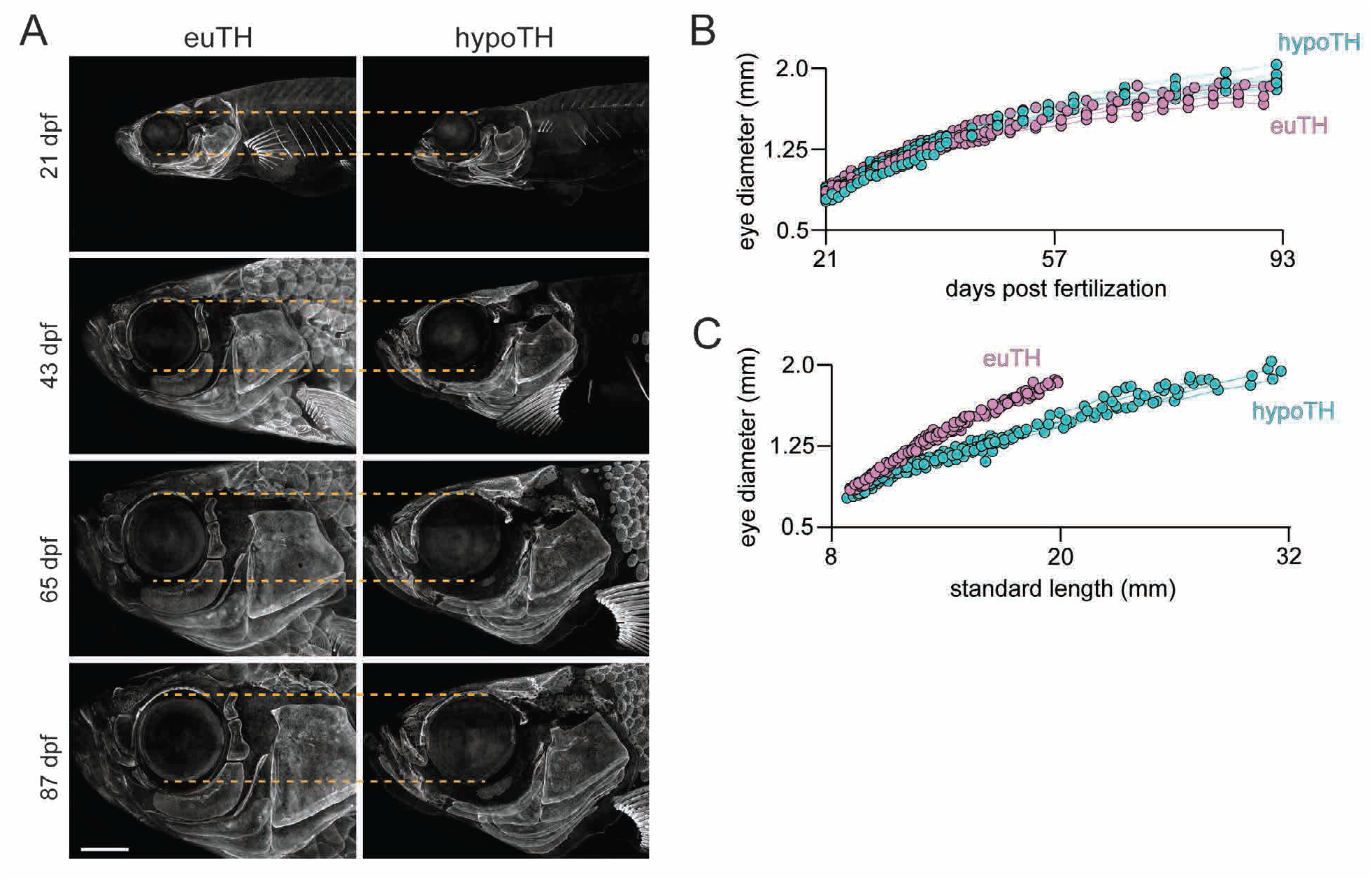
Eye growth was not affected by thyroid status. (**A**) Representative images from *sp7:EGFP* timeseries illustrating normal growth of the eye in hypoTH fish (orange dashed line), consistent with the observation that eyes are proportionally larger relative to body size in hypoTH fish compared to euTH at multiple stages of development (Hu et al., 2019). (**B**) Eye diameter increased at a similar rate in euTH and hypoTH fish (*n*=6 euTH; *n*=5 hypoTH). (**C**) Plot of eye diameter relative to standard length showing that the eye grew proportionally faster than the body in hypoTH fish compaed to euTH fish. Scale bar, (A) 1 mm.

**Fig. S3.**
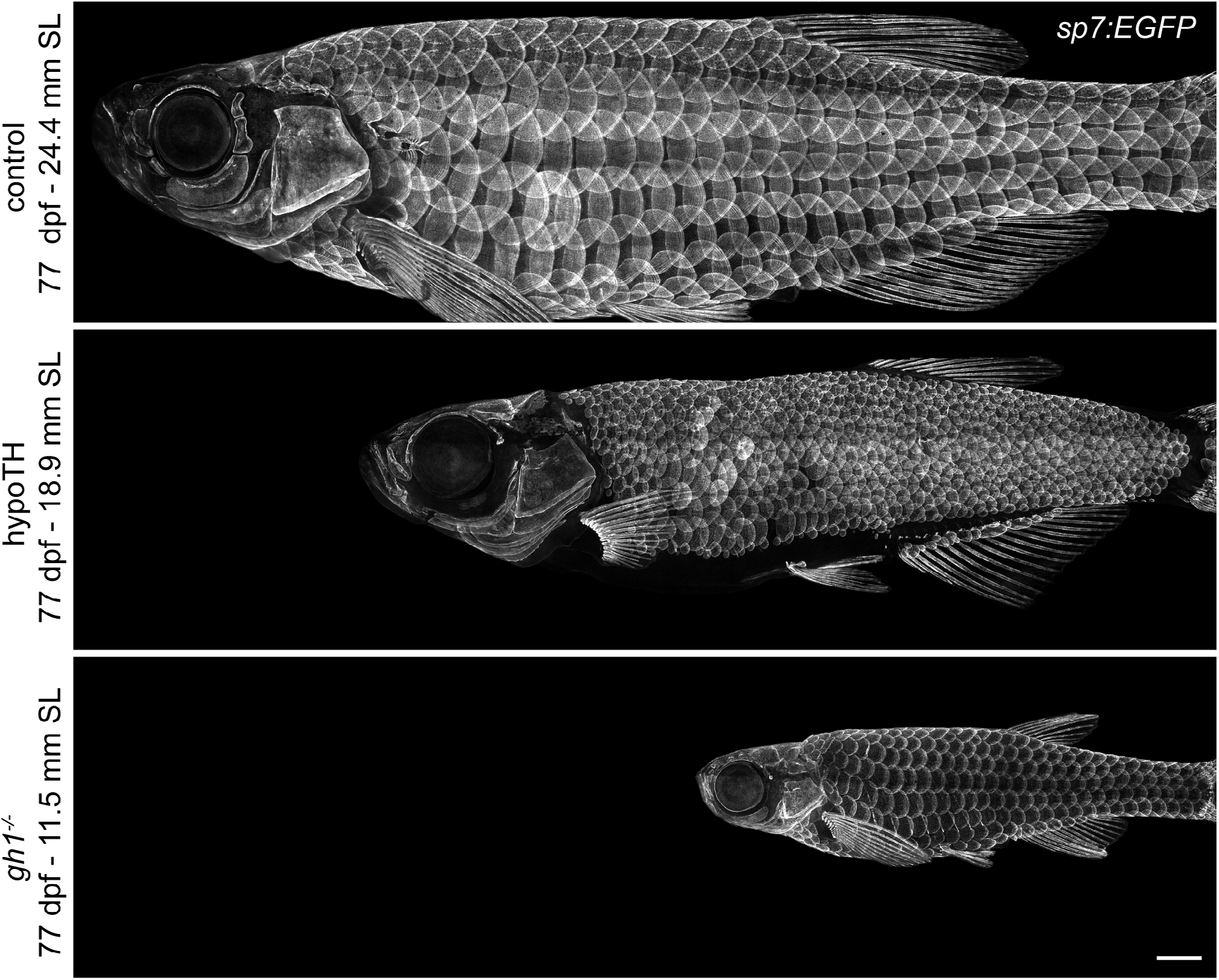
*gh1* mutants are smaller than euTH and hypoTH individuals as they approach adulthood. *sp7:EGFP* expressing euTH control, hypoTH and *gh1* mutants displayed to scale. The mutant is tiny yet had a normal pattern of squamation. Scale bar, 1 mm.

**Fig. S4.**
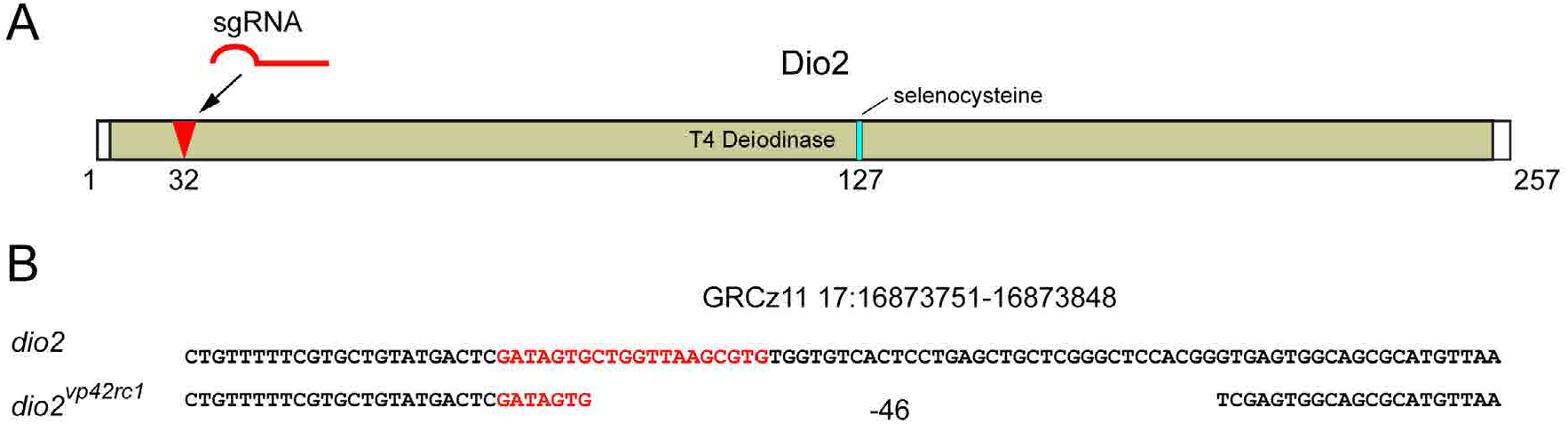
Generation of a *dio2* mutant allele. (**A**) Diagram of Dio2 protein sequence showing location of sgRNA target site (red arrow) within the deiodinase domain. Numbers indicate amino acid position relative to start codon. (**B**) Genomic location and genetic lesion of the *dio2* mutant, which leads to a frameshift followed by 6 novel amino acids and a premature stop codon. sgRNA target sequence indicated with red text.

**Fig. S5.**
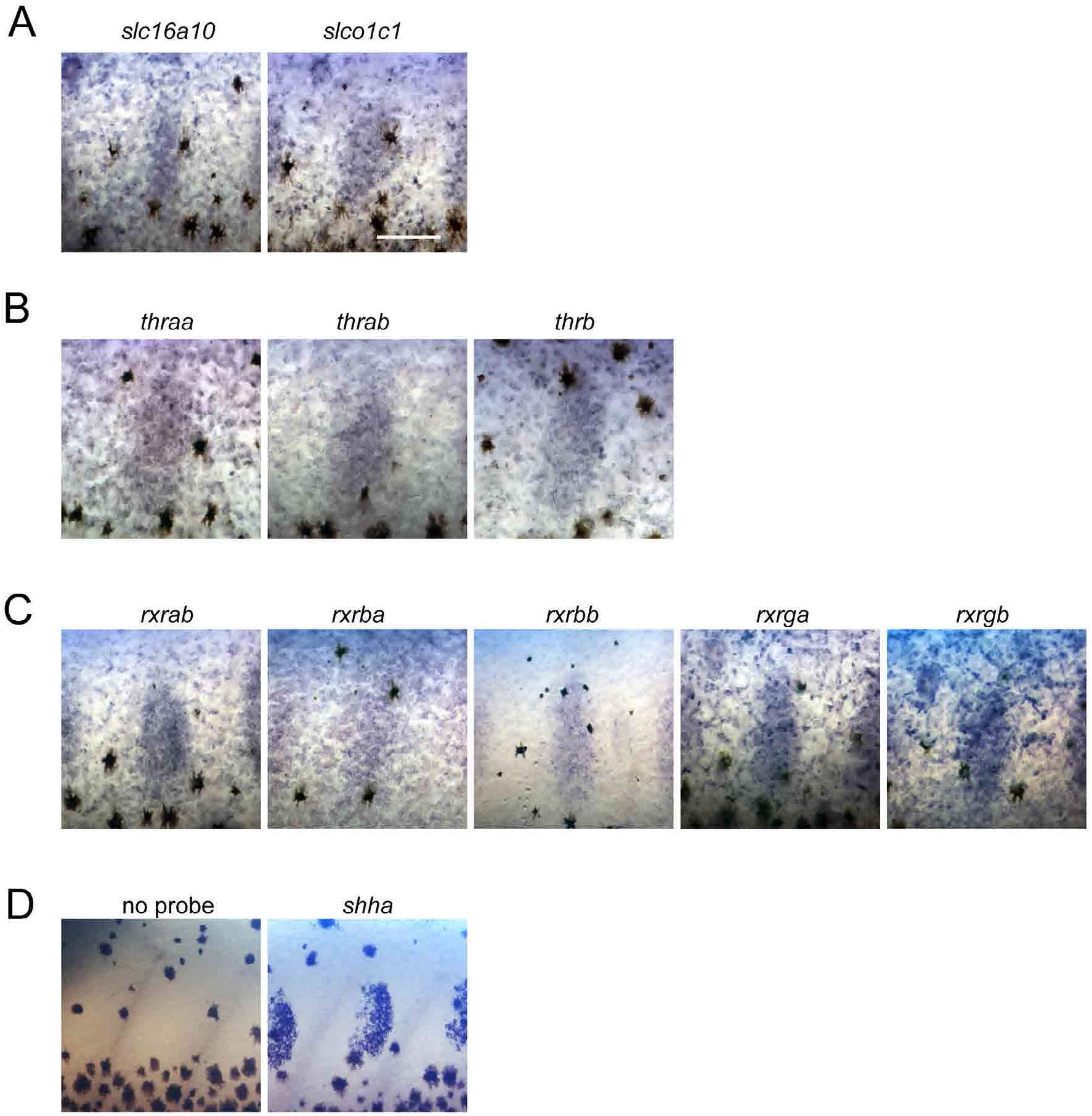
Expression TH pathway genes in skin during the period squamation. *In-situ* hybridization revealed transcripts for TH transporters (**A**), receptors (**B**) and co-receptors (**C**) in skin and scale papillae (one papilla is centered in each frame). (**E**) No-probe negative control; *shha* positive control. Scale bar, (in A for A–D) 200 μm.

**Fig. S6.**
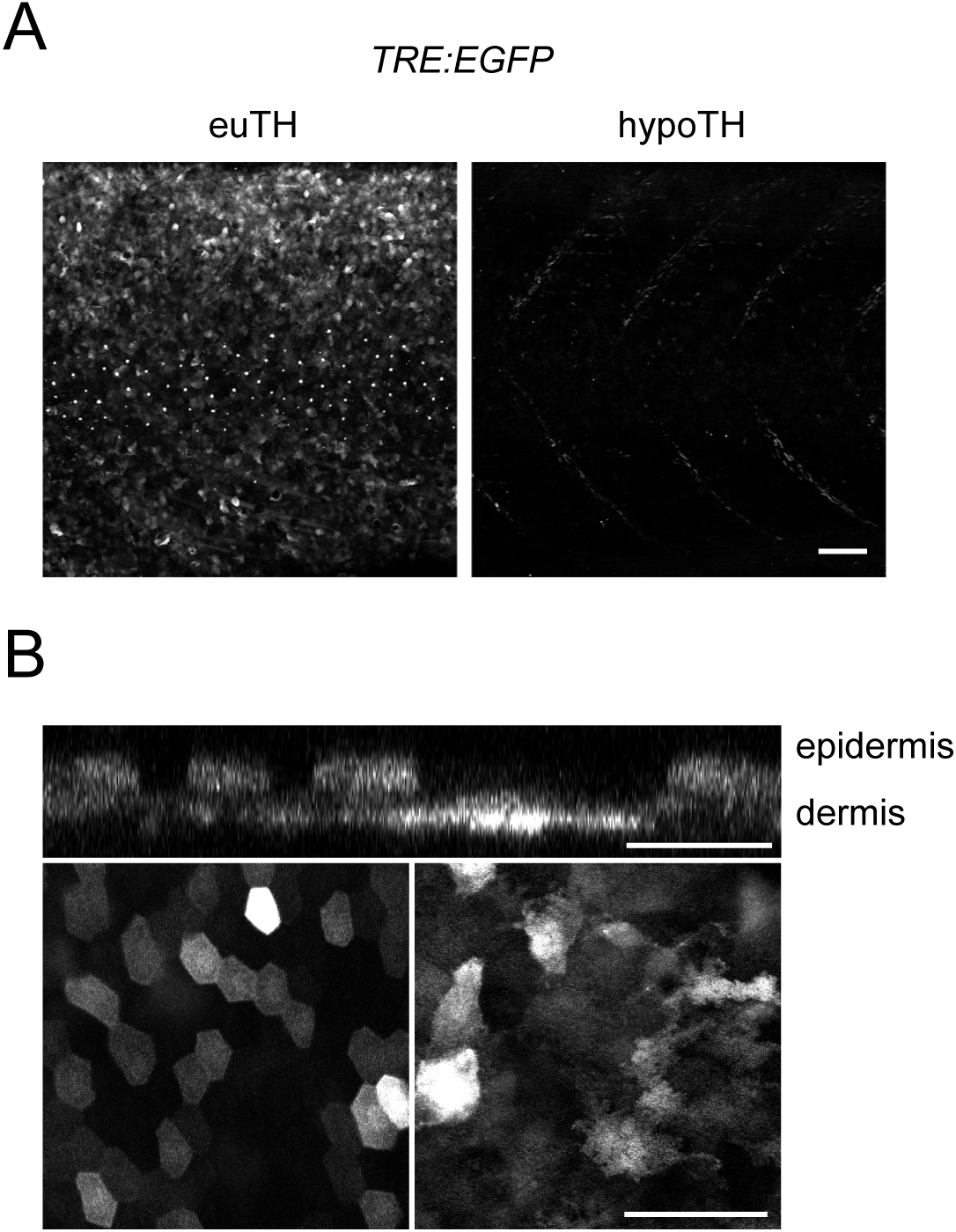
Expression of synthetic TH activity reporter in skin. (**A**)The synthetic TH reporter *TRE-bglob:EGFP* was broadly expressed in skin of euTH, but not hypoTH individuals (*n*=8 each condition). Weak fluorescence evident along the deeper horizontal myosepta in hypoTH represents TH independent expression of the reporter. (**B**) Higher magnification images showing reporter expression in superficial epidermis and deeper dermis in original planes of image capture (lower left and right, respectively) and in an orthogonal projection (above). All images were captured in 8.6 SSL larvae just prior to onset of squamation and before scales are overtly visible (Aman et al., 2018; Parichy et al., 2009). Scale bars, 100 μm.

**Fig. S7.**
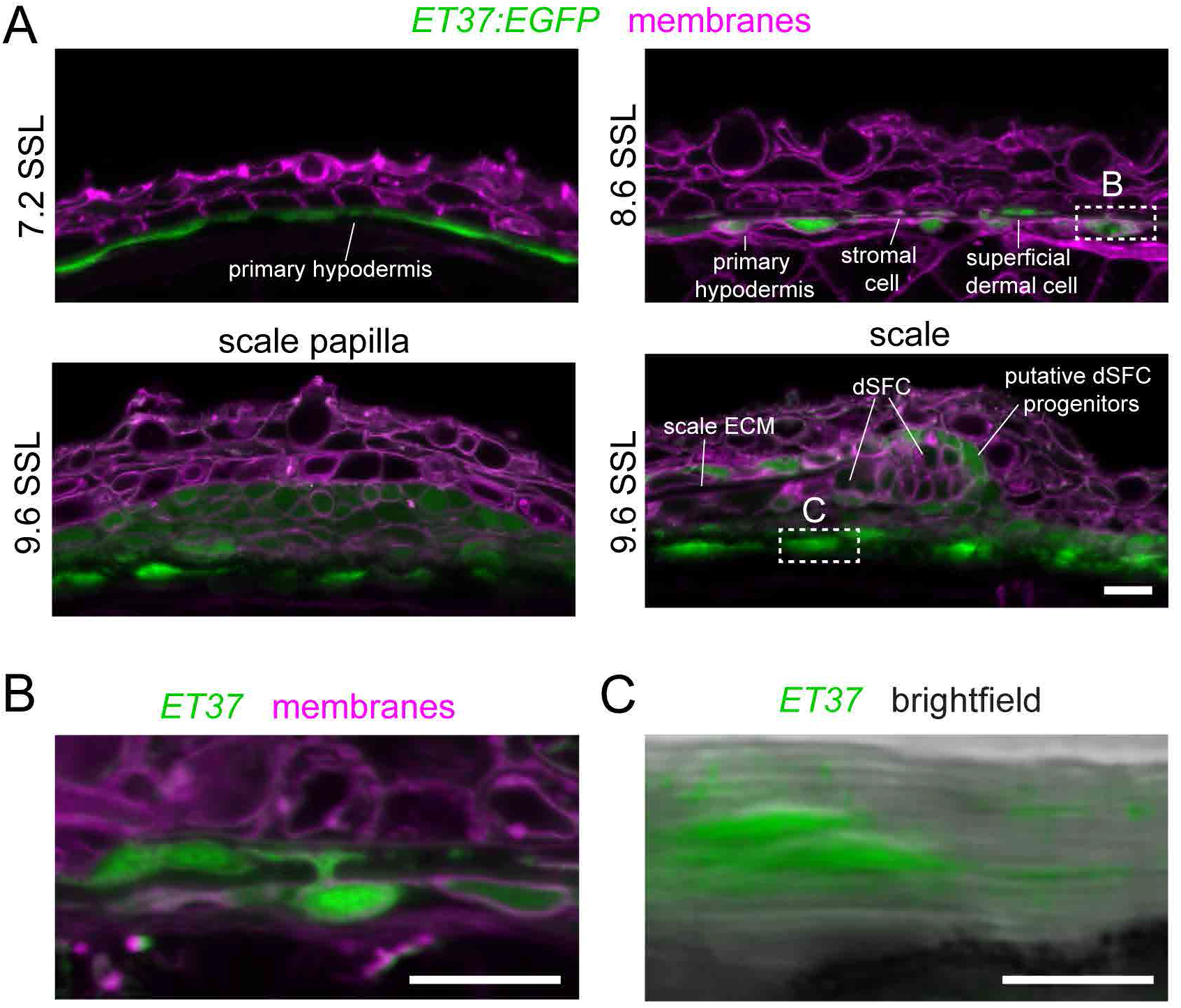
*ET37:EGFP* labels all dermal cells. Confocal images of vibratome sections through skin of *ET37:EGFP* transgenic larvae, counterstained with Cellbright membrane dye. (**A**) At 7.2 SSL, EGFP was expressed in a monolayer of cells beneath the epidermis, representing the primary hypodermis. At 8.6 SSL, the primary hypodermis was becoming a multilayered dermis. At 9.6 SSL, scale papillae and extending scales were present. *ET37:EGFP* was expressed in all papillar cells and in a subset of cells at the posterior margin of the scale. (**B**) Dermal cells labeled by *ET37:EGFP* included cells with projections extending from the cell body in a deep stratum of the developing dermis into a more superficial stratum towards the epidermis, as exemplified in this higher magnification image of the outlined region in A at 8.6 SSL. (**C**) *ET37:EGFP+* cells beneath scales were embedded in a thick collagenous stroma, as revealed by contrasting horizontal lines associated with fibrillar collagen in merged brightfield and fluorescence channels of the boxed region in A at 9.6 SSL. Each image is representative of sections from at least four individuals. Scale bars, 100 μm.

**Fig. S8.**
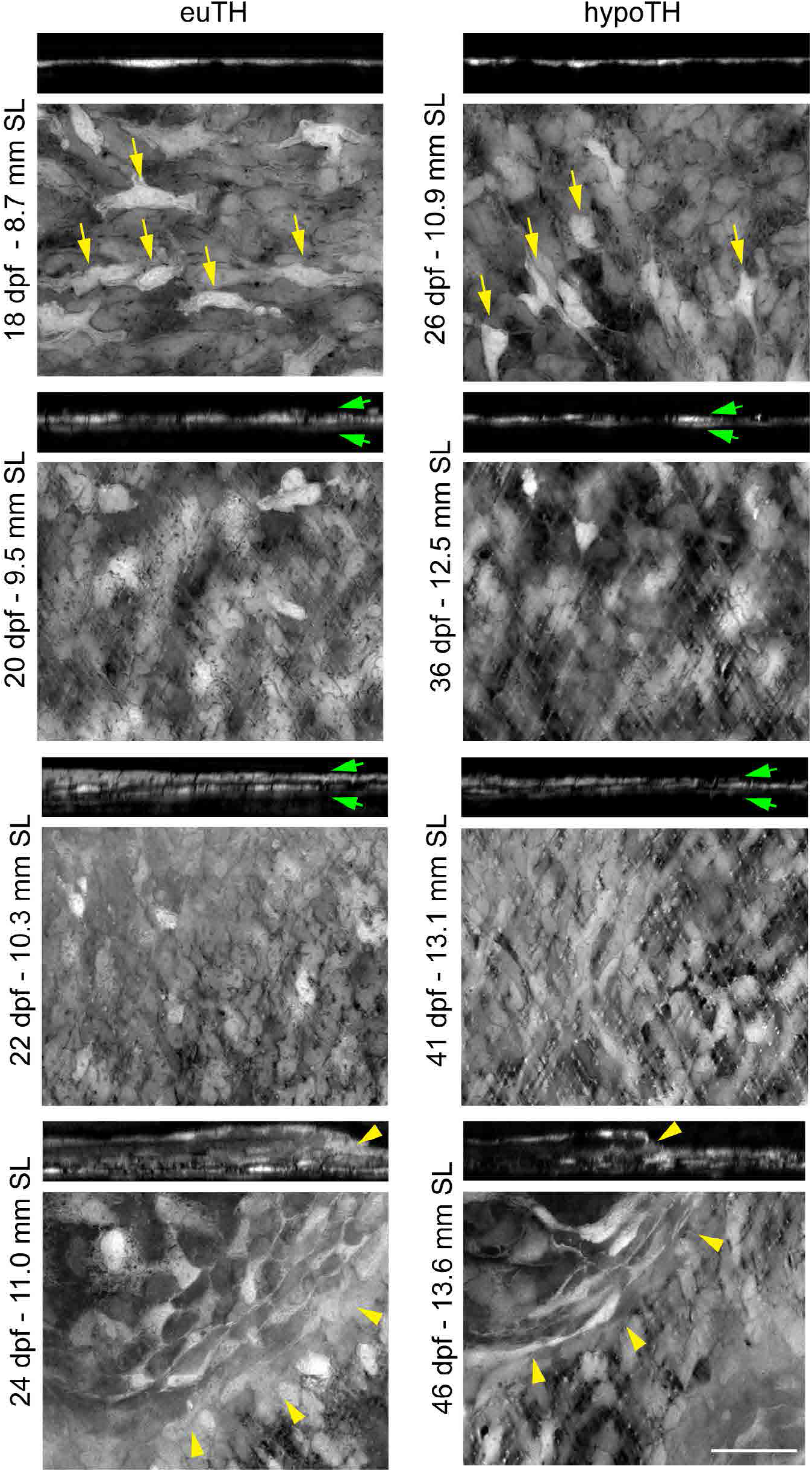
Dermal stratification followed a similar sequence of events in euTH and hypoTH individuals. Superresolution imaging showing the appearance of *ET37:EGFP* pre-migratory cells (yellow arrows), followed by stratified dermis (green arrows) and scales (yellow arrowheads). Orthogonal projection (top) and maximum intensity projection (bottom). Scale bar, 25 μm.

**Fig. S9.**
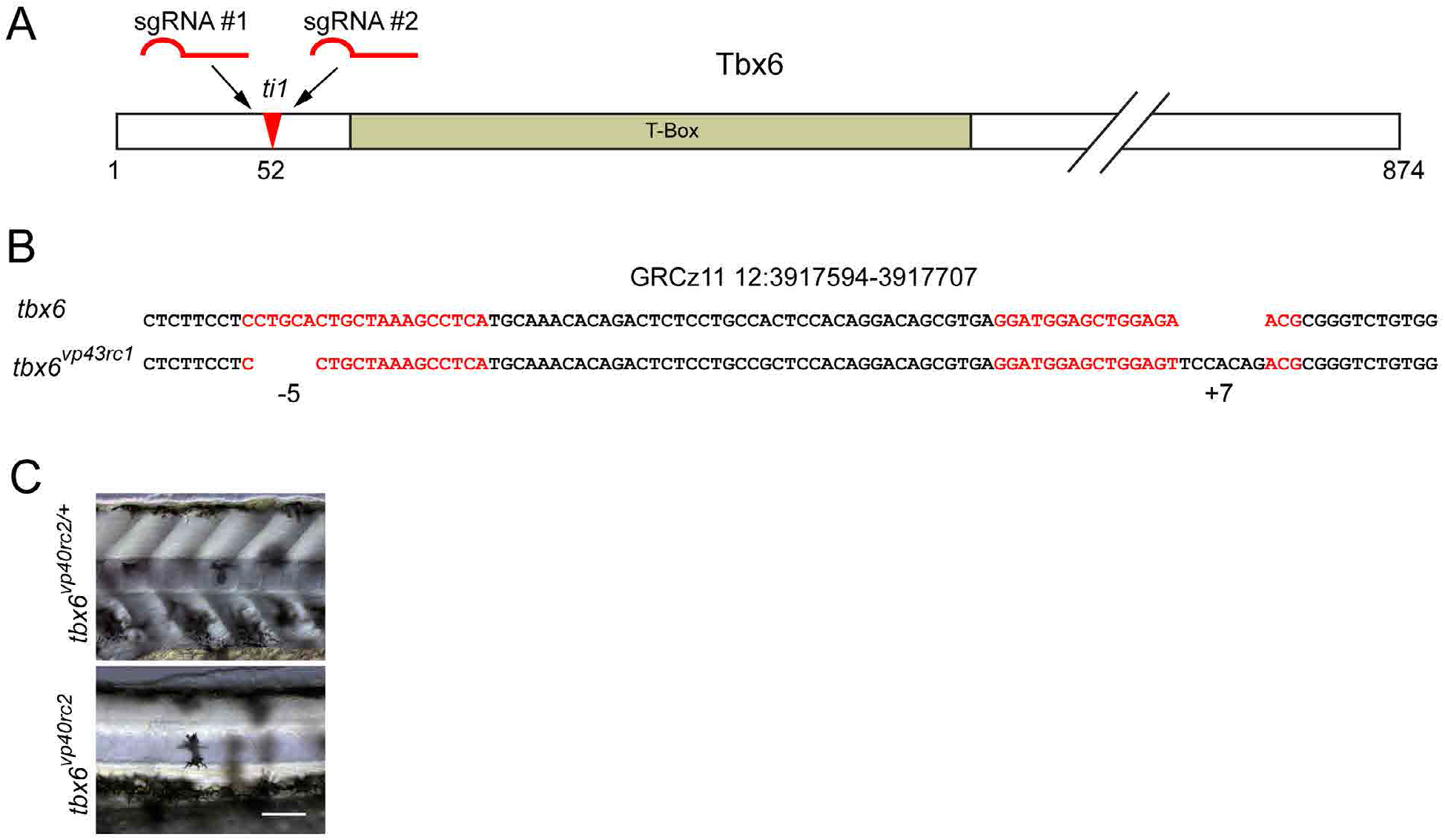
Generation of *tbx6* mutant allele. (**A**) Diagram of Tbx6 protein sequence showing location of sgRNA target sites in relation to the T-Box DNA binding domain and the previously reported *ti1* allele. (**B**) Genomic location and genetic lesion of the *tbx6* mutant. sgRNA target sequences indicated with red text. (**C**) At two days post fertilization, *tbx6* mutants displayed the classic fused-somites phenotype and lacked visible somites. Scale bar, 100 μm.

**Fig. S10.**
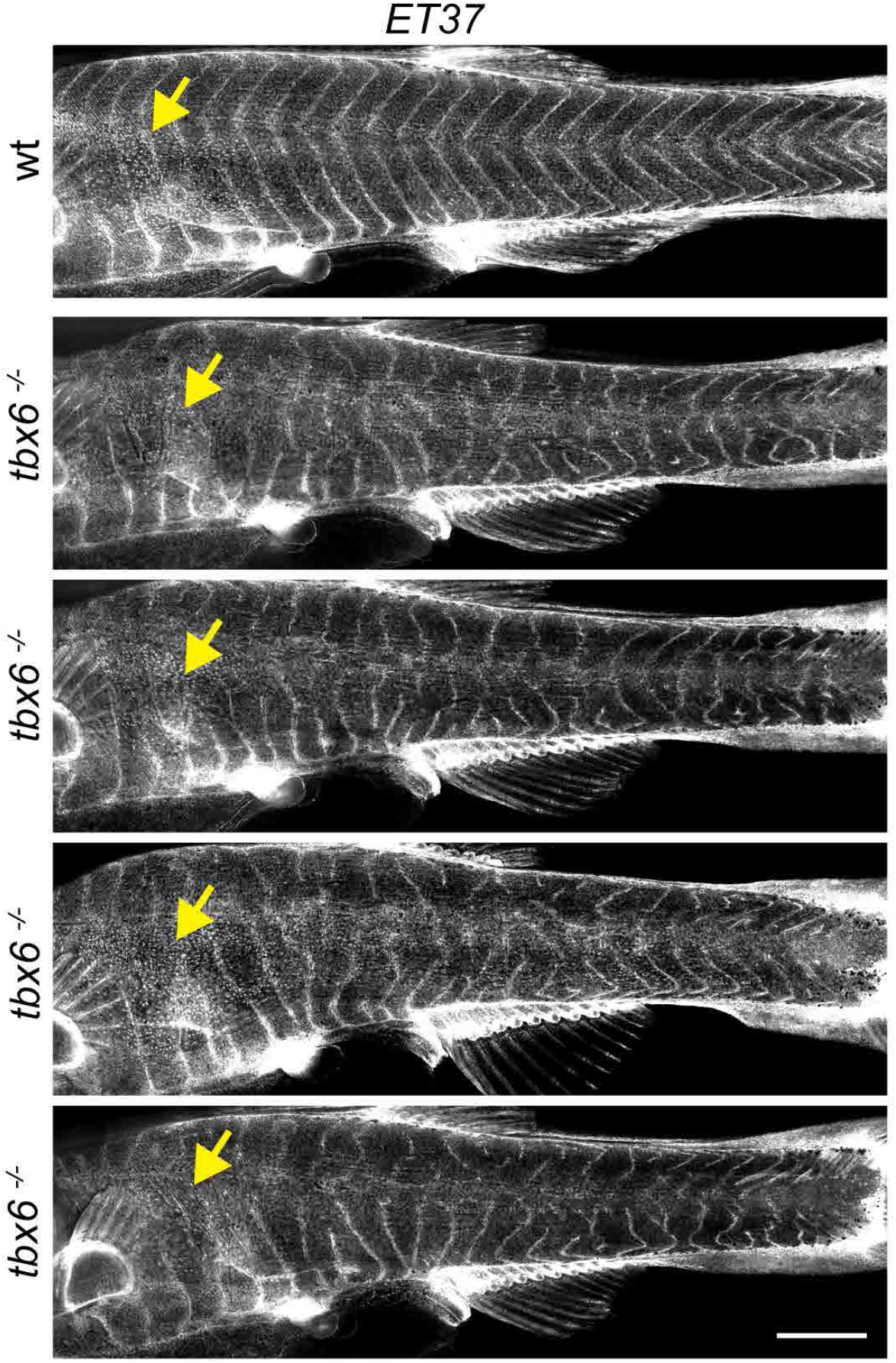
Segmentation in post-embryonic *tbx6* mutants. At 8.6 SSL, dermal cells were beginning to stratify (yellow arrows). *ET37:EGFP* expression in primary hypodermal cells that appeared to ensheath myotomes showed regularly spaced vertical myosepta in wild-type *tbx6* siblings. In mutants lacking somites during embryonic development, vertical myosepta were highly irregular. Shown are four representative examples of 12 fish. Scale bar, 1 mm.

**Fig. S11.**
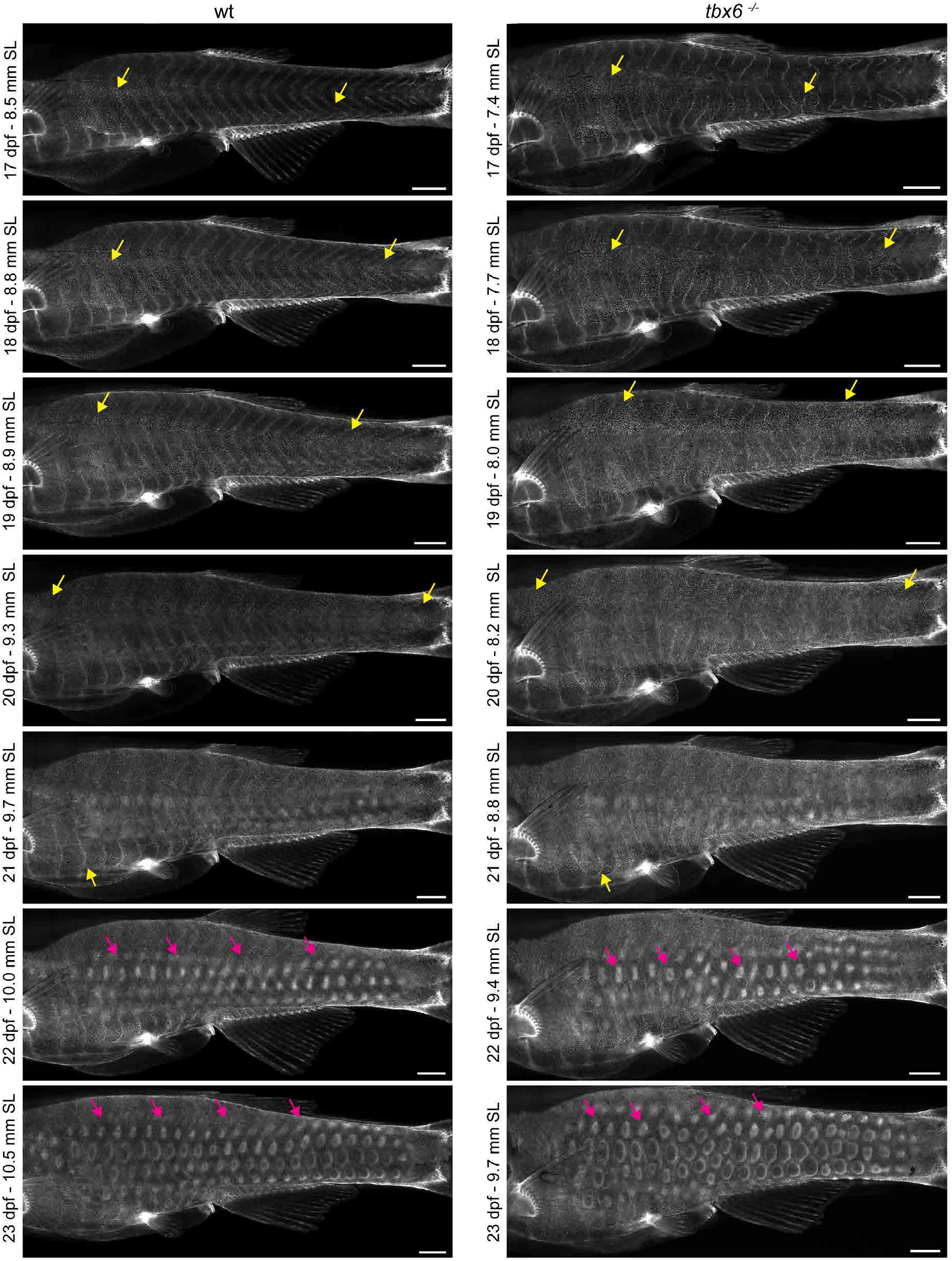
Dermal stratification in *tbx6* mutants. Repeated *in-toto* imaging of *ET37:EGFP* revealed that dermis in *tbx6* mutants stratified with patterning and timing like control animals. Yellow arrows indicate migrating primary dermal cells. Magenta arrows indicate developing scale papillae. Scale bars, 500 μm.

**Movie 1. Larval to adult transition in euTH and hypoTH fish registered to same image size.** *sp7:EGFP* expression in representative control (euTH) and hypoTH individuals shows squamation and relative growth of other elements of the dermal skeleton including the head skeleton and fin rays. Scale bar, 1 mm. Days post fertilization (DPF) provided as reference tempo.

**Movie 2. Larval to adult transition in euTH and hypoTH fish displayed to scale.** *sp7:EGFP* expression in representative control (euTH) and hypoTH displayed to scale to highlight differences in absolute growth rates. Scale bar, 1 mm. Days post fertilization (DPF) provided as reference tempo.

## References

Aman, A.J., Fulbright, A.N., Parichy, D.M., 2018. Wnt/beta-catenin regulates an ancient signaling network during zebrafish scale development. Elife 7.

Aman, A.J., Parichy, D.M., 2020. Chapter 8 - Zebrafish Integumentary System, in: Cartner, S.C., Eisen, J.S., Farmer, S.C., Guillemin, K.J., Kent, M.L., Sanders, G.E. (Eds.), The Zebrafish in Biomedical Research. Academic Press, pp. 91–96.

Bergh, J.J., Lin, H.-Y., Lansing, L., Mohamed, S.N., Davis, F.B., Mousa, S., Davis, P.J., 2005. Integrin αVβ3 Contains a Cell Surface Receptor Site for Thyroid Hormone that Is Linked to Activation of Mitogen-Activated Protein Kinase and Induction of Angiogenesis. Endocrinology 146, 2864–2871.

Bianco, A.C., Kim, B.W., 2006. Deiodinases: implications of the local control of thyroid hormone action. The Journal of clinical investigation 116, 2571–2579.

Bolotovskiy, A.A., Levina, M.A., DeFaveri, J., Merilä, J., Levin, B.A., 2018. Heterochronic development of lateral plates in the three-spined stickleback induced by thyroid hormone level alterations. PLOS ONE 13, e0194040.

Brent, G.A., 2012. Mechanisms of thyroid hormone action. The Journal of Clinical Investigation 122, 3035–3043.

Brown, D.D., Cai, L., 2007. Amphibian metamorphosis. Dev Biol 306, 20–33.

Buchholz, D.R., 2017. Xenopus metamorphosis as a model to study thyroid hormone receptor function during vertebrate developmental transitions. Mol Cell Endocrinol 459, 64–70.

Buchholz, D.R., Hsia, S.-C.V., Fu, L., Shi, Y.-B., 2003. A Dominant-Negative Thyroid Hormone Receptor Blocks Amphibian Metamorphosis by Retaining Corepressors at Target Genes. Molecular and Cellular Biology 23, 6750–6758.

Cheng, C.W., Niu, B., Warren, M., Pevny, L.H., Lovell-Badge, R., Hwa, T., Cheah, K.S.E., 2014. Predicting the spatiotemporal dynamics of hair follicle patterns in the developing mouse. Proceedings of the National Academy of Sciences 111, 2596–2601.

Concordet, J.-P., Haeussler, M., 2018. CRISPOR: intuitive guide selection for CRISPR/Cas9 genome editing experiments and screens. Nucleic Acids Research 46, W242–W245.

Cooper, R.L., Thiery, A.P., Fletcher, A.G., Delbarre, D.J., Rasch, L.J., Fraser, G.J., 2018. An ancient Turing-like patterning mechanism regulates skin denticle development in sharks. Sci. Adv. 4, eaau5484.

Cox, B.D., De Simone, A., Tornini, V.A., Singh, S.P., Di Talia, S., Poss, K.D., 2018. In Toto Imaging of Dynamic Osteoblast Behaviors in Regenerating Skeletal Bone. Curr Biol 28, 3937–3947 e3934.

Dalle Nogare, D., Chitnis, A.B., 2017. Self-organizing spots get under your skin. PLOS Biology 15, e2004412.

Davis, P.J., Davis, F.B., 1996. Nongenomic actions of thyroid hormone. Thyroid: official journal of the American Thyroid Association 6, 497–504.

De Simone, A., Evanitsky, M.N., Hayden, L., Cox, B.D., Wang, J., Tornini, V.A., Ou, J., Chao, A., Poss, K.D., Di Talia, S., 2021. Control of osteoblast regeneration by a train of Erk activity waves. Nature.

DeLaurier, A., Eames, B.F., Blanco-Sanchez, B., Peng, G., He, X., Swartz, M.E., Ullmann, B., Westerfield, M., Kimmel, C.B., 2010. Zebrafish sp7:EGFP: a transgenic for studying otic vesicle formation, skeletogenesis, and bone regeneration. Genesis 48, 505–511.

Eom, D.S., Patterson, L.B., Bostic, R.R., Parichy, D.M., 2021. Immunoglobulin superfamily receptor Junctional adhesion molecule 3 (Jam3) requirement for melanophore survival and patterning during formation of zebrafish stripes. bioRxiv, 2021.2003.2001.433381.

Goodrich, H.B., Nichols, R., 1933. SCALE TRANSPLANTATION IN THE GOLDFISH CARASSIUS AURATUS: I. EFFECTS ON CHROMATOPHORES. II. TISSUE REACTIONS. The Biological Bulletin 65, 253–265.

Green, J.B., Sharpe, J., 2015. Positional information and reaction-diffusion: two big ideas in developmental biology combine. Development 142, 1203–1211.

Hammes, S.R., Davis, P.J., 2015. Overlapping nongenomic and genomic actions of thyroid hormone and steroids. Best Practice & Research Clinical Endocrinology & Metabolism 29, 581–593.

Han, C.R., Holmsen, E., Carrington, B., Bishop, K., Zhu, Y.J., Starost, M., Meltzer, P., Sood, R., Liu, P., Cheng, S.Y., 2020. Generation of Novel Genetic Models to Dissect Resistance to Thyroid Hormone Receptor α in Zebrafish. Thyroid: official journal of the American Thyroid Association 30, 314–328.

Harris, M.P., Rohner, N., Schwarz, H., Perathoner, S., Konstantinidis, P., Nüsslein-Volhard, C., 2008. Zebrafish *eda* and *edar* Mutants Reveal Conserved and Ancestral Roles of Ectodysplasin Signaling in Vertebrates. PLoS Genet 4, e1000206.

Hiscock, T.W., Megason, S.G., 2015. Mathematically guided approaches to distinguish models of periodic patterning. Development 142, 409–419.

Ho, W.K.W., Freem, L., Zhao, D., Painter, K.J., Woolley, T.E., Gaffney, E.A., McGrew, M.J., Tzika, A., Milinkovitch, M.C., Schneider, P., Drusko, A., Matthaus, F., Glover, J.D., Wells, K.L., Johansson, J.A., Davey, M.G., Sang, H.M., Clinton, M., Headon, D.J., 2019. Feather arrays are patterned by interacting signalling and cell density waves. PLoS Biol 17, e3000132.

Hörlein, A.J., Näär, A.M., Heinzel, T., Torchia, J., Gloss, B., Kurokawa, R., Ryan, A., Kamei, Y., Söderström, M., Glass, C.K., Rosenfeld, M.G., 1995. Ligand-independent repression by the thyroid hormone receptor mediated by a nuclear receptor co-repressor. Nature 377, 397–404.

Hu, Y., Harper, M., Acosta, B., Donahue, J., Bui, H., Lee, H., Nguyen, S., McMenamin, S., 2020. Thyroid hormone regulates proximodistal identity in the fin skeleton. bioRxiv, 2020.2008.2018.256354.

Hu, Y., Mauri, A., Donahue, J., Singh, R., Acosta, B., McMenamin, S., 2019. Thyroid hormone coordinates developmental trajectories but does not underlie developmental truncation in danionins. Dev Dyn 248, 1144–1154.

Hur, M., Gistelinck, C.A., Huber, P., Lee, J., Thompson, M.H., Monstad-Rios, A.T., Watson, C.J., McMenamin, S.K., Willaert, A., Parichy, D.M., Coucke, P., Kwon, R.Y., 2017. MicroCT-based phenomics in the zebrafish skeleton reveals virtues of deep phenotyping in a distributed organ system. eLife 6, e26014.

Iglesias, J., Rodríguez-Ojea, G., 1997. The use of alizarin complexone for immersion marking of the otoliths of embryos and larvae of the turbot, Scophthalmus maximus (L.): dosage and treatment time. Fish Manage Ecol 4.

Iwasaki, M., Kuroda, J., Kawakami, K., Wada, H., 2018. Epidermal regulation of bone morphogenesis through the development and regeneration of osteoblasts in the zebrafish scale. Dev Biol.

Kim, H.-Y., Mohan, S., 2013. Role and Mechanisms of Actions of Thyroid Hormone on the Skeletal Development. Bone Research 1, 146–161.

Kulin, H.E., Müller, J., 1996. The Biological Aspects of Puberty. Pediatrics in Review 17, 75–86.

Kumar, M., Sarma, D.K., Shubham, S., Kumawat, M., Verma, V., Prakash, A., Tiwari, R., 2020. Environmental Endocrine-Disrupting Chemical Exposure: Role in Non-Communicable Diseases. Frontiers in Public Health 8.

Lai, Y.C., Chuong, C.M., 2016. The “tao” of integuments. Science 354, 1533–1534.

Laudet, V., 2011. The Origins and Evolution of Vertebrate Metamorphosis. Current Biology 21, R726–R737.

Le Guellec, D., Morvan-Dubois, G., Sire, J.Y., 2004. Skin development in bony fish with particular emphasis on collagen deposition in the dermis of the zebrafish (Danio rerio). Int J Dev Biol 48, 217–231.

Levin, B.A., 2010. Drastic shift in the number of lateral line scales in the common roach Rutilus rutilus as a result of heterochronies: experimental data. Journal of Applied Ichthyology 26, 303–306.

Mancino, G., Miro, C., Di Cicco, E., Dentice, M., 2021. Thyroid hormone action in epidermal development and homeostasis and its implications in the pathophysiology of the skin. Journal of Endocrinological Investigation.

Matsuda, H., 2018. Zebrafish as a model for studying functional pancreatic β cells development and regeneration. Dev Growth Differ 60, 393–399.

Matsuda, H., Mullapudi, S.T., Zhang, Y., Hesselson, D., Stainier, D.Y.R., 2017. Thyroid Hormone Coordinates Pancreatic Islet Maturation During the Zebrafish Larval-to-Juvenile Transition to Maintain Glucose Homeostasis. Diabetes 66, 2623–2635.

McMenamin, S.K., Bain, E.J., McCann, A.E., Patterson, L.B., Eom, D.S., Waller, Z.P., Hamill, J.C., Kuhlman, J.A., Eisen, J.S., Parichy, D.M., 2014. Thyroid hormone-dependent adult pigment cell lineage and pattern in zebrafish. Science 345, 1358–1361.

McMenamin, S.K., Minchin, J.E.N., Gordon, T.N., Rawls, J.F., Parichy, D.M., 2013. Dwarfism and Increased Adiposity in the gh1 Mutant Zebrafish vizzini. Endocrinology 154, 1476–1487.

McMenamin, S.K., Parichy, D.M., 2013. Metamorphosis in teleosts. Current topics in developmental biology 103, 127–165.

Moro, E., Ozhan-Kizil, G., Mongera, A., Beis, D., Wierzbicki, C., Young, R.M., Bournele, D., Domenichini, A., Valdivia, L.E., Lum, L., Chen, C., Amatruda, J.F., Tiso, N., Weidinger, G., Argenton, F., 2012. In vivo Wnt signaling tracing through a transgenic biosensor fish reveals novel activity domains. Developmental Biology 366, 327–340.

Morris, J.L., Cross, S.J., Lu, Y., Kadler, K.E., Lu, Y., Dallas, S.L., Martin, P., 2018. Live imaging of collagen deposition during skin development and repair in a collagen I - GFP fusion transgenic zebrafish line. Developmental biology 441, 4–11.

Mosimann, C., Kaufman, C.K., Li, P., Pugach, E.K., Tamplin, O.J., Zon, L.I., 2011. Ubiquitous transgene expression and Cre-based recombination driven by the ubiquitin promoter in zebrafish. Development 138, 169–177.

Nikaido, M., Kawakami, A., Sawada, A., Furutani-Seiki, M., Takeda, H., Araki, K., 2002. Tbx24, encoding a T-box protein, is mutated in the zebrafish somite-segmentation mutant fused somites. Nat Genet 31, 195–199.

Ouyang, Q., Swinney, H.L., 1991. Transition from a uniform state to hexagonal and striped Turing patterns. Nature 352, 610–612.

Painter, K.J., Hunt, G.S., Wells, K.L., Johansson, J.A., Headon, D.J., 2012. Towards an integrated experimental–theoretical approach for assessing the mechanistic basis of hair and feather morphogenesis. Interface focus 2, 433–450.

Pan, Y.A., Freundlich, T., Weissman, T.A., Schoppik, D., Wang, X.C., Zimmerman, S., Ciruna, B., Sanes, J.R., Lichtman, J.W., Schier, A.F., 2013. Zebrabow: multispectral cell labeling for cell tracing and lineage analysis in zebrafish. Development 140, 2835–2846.

Parichy, D.M., Elizondo, M.R., Mills, M.G., Gordon, T.N., Engeszer, R.E., 2009. Normal table of postembryonic zebrafish development: Staging by externally visible anatomy of the living fish. Developmental Dynamics 238, 2975–3015.

Parinov, S., Kondrichin, I., Korzh, V., Emelyanov, A., 2004. Tol2 transposon-mediated enhancer trap to identify developmentally regulated zebrafish genes in vivo. Dev Dyn 231, 449–459.

Pyati, U.J., Webb, A.E., Kimelman, D., 2005. Transgenic zebrafish reveal stage-specific roles for Bmp signaling in ventral and posterior mesoderm development. Development 132, 2333–2343.

Quigley, I.K., Turner, J.M., Nuckels, R.J., Manuel, J.L., Budi, E.H., MacDonald, E.L., Parichy, D.M., 2004. Pigment pattern evolution by differential deployment of neural crest and post-embryonic melanophore lineages in Danio fishes. Development 131, 6053–6069.

Rasmussen, J.P., Vo, N.T., Sagasti, A., 2018. Fish Scales Dictate the Pattern of Adult Skin Innervation and Vascularization. Dev Cell 46, 344–359 e344.

Renfer, E., Amon-Hassenzahl, A., Steinmetz, P., Technau, U., 2009. A Muscle-Specific Transgenic Reporter Line of the Sea Anemone, Nematostella Vectensis. Proceedings of the National Academy of Sciences of the United States of America 107, 104–108.

Renn, J., Winkler, C., 2009. Osterix-mCherry transgenic medaka for vivo imaging of bone formation. Developmental dynamics: an official publication of the American Association of Anatomists 238, 241–248.

Rohner, N., Bercsényi, M., Orbán, L., Kolanczyk, M.E., Linke, D., Brand, M., Nüsslein-Volhard, C., Harris, M.P., 2009. Duplication of fgfr1 Permits Fgf Signaling to Serve as a Target for Selection during Domestication. Current Biology 19, 1642–1647.

Saunders, L.M., Mishra, A.K., Aman, A.J., Lewis, V.M., Toomey, M.B., Packer, J.S., Qiu, X., McFaline-Figueroa, J.L., Corbo, J.C., Trapnell, C., Parichy, D.M., 2019. Thyroid hormone regulates distinct paths to maturation in pigment cell lineages. eLife 8, e45181.

Schindelin, J., Arganda-Carreras, I., Frise, E., Kaynig, V., Longair, M., Pietzsch, T., Preibisch, S., Rueden, C., Saalfeld, S., Schmid, B., Tinevez, J.-Y., White, D.J., Hartenstein, V., Eliceiri, K., Tomancak, P., Cardona, A., 2012. Fiji: an open-source platform for biological-image analysis. Nature Methods 9, 676–682.

Shah, A.N., Davey, C.F., Whitebirch, A.C., Miller, A.C., Moens, C.B., 2015. Rapid reverse genetic screening using CRISPR in zebrafish. Nature Methods 12, 535–540.

Shi, Y.B., 2000. Amphibian Metamorphosis: From Morphology to Molecular Biology. John Wiley and Sons, Inc., New York, New York.

Sire, J.-Y., Allizard, F., Babiar, O., Bourguignon, J., Quilhac, A., 1997. Scale development in zebrafish (Danio rerio). Journal of Anatomy 190, 545–561.

Sire, J.Y., Akimenko, M.A., 2004. Scale development in fish: a review, with description of sonic hedgehog (shh) expression in the zebrafish (Danio rerio). Int J Dev Biol 48, 233–247.

Skah, S., Uchuya-Castillo, J., Sirakov, M., Plateroti, M., 2017. The thyroid hormone nuclear receptors and the Wnt/β-catenin pathway: An intriguing liaison. Developmental Biology 422, 71–82.

Slootweg, M.C., 1993. Growth hormone and bone. Hormone and metabolic research = Hormon- und Stoffwechselforschung = Hormones et metabolisme 25, 335–343.

Smirnov, S.V., Dzerzhinskii, K.F., Levin, B.A., 2006. On the relationship between scale number in the lateral line in the african barbel Barbus intermedius and the rate of ontogeny (by experimental data). Journal of Ichthyology 46, 129–132.

Sokal, R.R., Rohlf, F.J., 1981. Biometry. W. H. Freeman and Company, New York, New York.

Sosa-Ferrera, Z., Mahugo-Santana, C., Santana-Rodríguez, J.J., 2013. Analytical Methodologies for the Determination of Endocrine Disrupting Compounds in Biological and Environmental Samples. BioMed Research International 2013, 674838.

Tata, J.R., 1998. Amphibian metamorphosis as a model for studying the developmental actions of thyroid hormone. Cell Research 8, 259–272.

Temereva, E.N., Tsitrin, E.B., 2013. Development, organization, and remodeling of phoronid muscles from embryo to metamorphosis (Lophotrochozoa: Phoronida). BMC Dev Biol 13, 14.

van Eeden, F.J., Granato, M., Schach, U., Brand, M., Furutani-Seiki, M., Haffter, P., Hammerschmidt, M., Heisenberg, C.P., Jiang, Y.J., Kane, D.A., Kelsh, R.N., Mullins, M.C., Odenthal, J., Warga, R.M., Allende, M.L., Weinberg, E.S., Nusslein-Volhard, C., 1996. Mutations affecting somite formation and patterning in the zebrafish, Danio rerio. Development 123, 153164.

Volkov, L.I., Kim-Han, J.S., Saunders, L.M., Poria, D., Hughes, A.E.O., Kefalov, V.J., Parichy, D.M., Corbo, J.C., 2020. Thyroid hormone receptors mediate two distinct mechanisms of long-wavelength vision. Proceedings of the National Academy of Sciences 117, 15262–15269.

Widelitz, R.B., Jiang, T.-X., Noveen, A., Chen, C.-W.J., Chuong, C.-M., 1996. FGF Induces New Feather Buds From Developing Avian Skin. Journal of Investigative Dermatology 107, 797–803.

Windner, S.E., Bird, N.C., Patterson, S.E., Doris, R.A., Devoto, S.H., 2012. Fss/Tbx6 is required for central dermomyotome cell fate in zebrafish. Biology Open 1, 806–814.

Woltmann, I., Shkil, F., De Clercq, A., Huysseune, A., Witten, P.E., 2018. Supernumerary teeth in the pharyngeal dentition of slow-developing zebrafish (Danio rerio, Hamilton, 1822). Journal of Applied Ichthyology 34, 455–464.

Zhang, J., Wagh, P., Guay, D., Sanchez-Pulido, L., Padhi, B.K., Korzh, V., Andrade-Navarro, M.A., Akimenko, M.-A., 2010. Loss of fish actinotrichia proteins and the fin-to-limb transition. Nature 466, 234–237.

